# Song distinguishability predicts reproductive isolation between subspecies of the dark-eyed junco

**DOI:** 10.1101/2023.09.14.557432

**Authors:** Sarah Hourihan, Emily Hudson, Ximena León Du’Mottuchi, Emily Beach, Sydnie Smith, Nicole Creanza

## Abstract

The dark-eyed junco (*Junco hyemalis*) has experienced rapid phenotypic diversification within the last 18,000 years, resulting in several subspecies that reside in partially overlapping regions across North America. These subspecies have distinct plumage and morphology. If members of a subspecies disproportionately mate with one another, we would expect genetic differences to accumulate between the subspecies. In parallel, their learned songs could also accumulate changes. If song is used by individuals to recognize members of their own subspecies during mate selection, which would prevent the production of less fit hybrid offspring between subspecies, then song differences might co-localize with subspecies boundaries. Here, we quantify 10 song features to explore subspecies-level song variation using song recordings from community-science databases. We build a machine learning classifier to measure how accurately the subspecies’ songs can be distinguished from one another. Here, we show that songs of dark-eyed junco subspecies exhibit significant song-feature differences. However, these differences do not necessarily lead to distinguishability between subspecies. Notably, we find that subspecies pairs with adjacent ranges that do not hybridize have much more distinguishable songs, and also more evidence for genetic differentiation, than pairs that are known to hybridize. Thus, song distinguishability appears to have predictive power about which subspecies will hybridize, suggesting that song might play a role in reinforcing certain subspecies boundaries more than others. Finally, we analyze subspecies-level song differences alongside available genetic data and geographic coordinates to characterize the current evolutionary landscape of the dark-eyed junco subspecies complex. We observe geographic signal in the song and genetic data, indicating that individuals who share a range are more likely to share song characteristics and be genetically similar. This study illuminates the existence of subspecies-level song differences in the dark-eyed junco and provides further clarity on the role learned song plays in reinforcing reproductive boundaries between dark-eyed junco subspecies.

## INTRODUCTION

One of the most fundamental questions in biology is “How do new species form?” [1] A critical stage of species formation is when two groups of organisms become reproductively isolated from one another, such that they generally do not produce offspring [2]. In birds, the capability to hybridize between species appears to persist for millions of years [3]. Because hybridization between closely related species is physiologically possible yet rare [4–6], species boundaries in birds are thought to be maintained by pre-mating mechanisms, such as visual and auditory signals [7,8]. One auditory signal is birdsong, a set of vocalizations that function in species recognition, mate attraction, and territory defense [9,10]. In the oscine songbirds (Order: Passeriformes), birdsong is a learned behavior [11,12]; juveniles learn their species-typical songs from adult conspecific tutors. Thus, birdsong has notable potential to be a reproductive isolation mechanism that is learned. Consequently, birdsong could have the ability to accelerate speciation and diversification since culturally transmitted traits can evolve faster than genetically heritable ones [13,14].

To study the evolution of birdsong and its role in maintaining species boundaries, we focus on the dark-eyed junco (*Junco hyemalis*)—a common North American songbird that exhibits geographic variation spanning from local populations to subspecies, and from complete geographic isolation to regions of overlap with interbreeding forms [15–18].

The dark-eyed junco has diverse plumage and morphology within the species, with six major subspecies distributed across North America: the slate-colored junco (*J. hyemalis hyemalis*) in eastern and boreal North America, the white-winged junco (*J. hyemalis aikeni*) in the northern Great Plains of the United States, the Oregon junco (*J. hyemalis oreganus*) spanning the west coast from California to Alaska, the pink-sided junco (*J. hyemalis mearnsi*) in northwestern United States, the gray-headed junco (*J. hyemalis caniceps*) in the Rocky Mountains region, and the red-backed junco (*J. hyemalis dorsalis*) in southwestern United States [16] (**Fig. 1**). The geographic breeding ranges of the dark-eyed junco subspecies have a small degree of overlap at their boundaries, but sightings of putative between-subspecies hybrids account for less than 1% of total sightings in regions of overlap (**Fig. 1, Table S1, Supplemental Materials**), indicating that plumage distinctions between the subspecies are maintained even in overlapping breeding areas.

**Figure 1.**
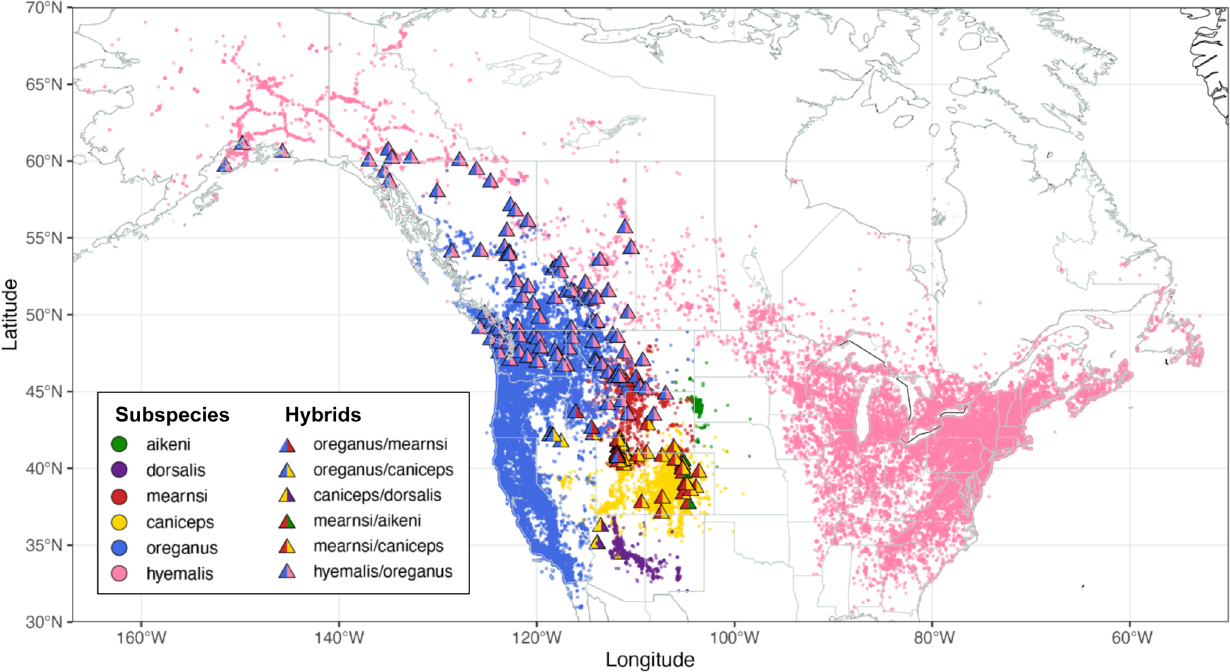
Breeding season distribution map for the dark-eyed junco. This map was constructed using observation data contributed to eBird and includes sightings of hybrid individuals (indicated by triangles).

Diversification of dark-eyed junco subspecies is an example of potential speciation in action within a rapidly evolving genus. Spatial genetic patterns of mitochondrial DNA (mtDNA) found that dark-eyed junco subspecies have genetic signatures of recent population expansion coinciding with the Last Glacial Maximum (∼18 ka) [19].

Plumage differences and genetic differentiation suggest that these subspecies might not interbreed very often, raising the question of whether song differences have also emerged between the subspecies, which no studies have yet assessed. If the subspecies’ songs are found to be distinct, we hypothesize that song plays a role in maintaining subspecies boundaries [20–23], in addition to visual plumage differences and geographic separation. For subspecies pairs that have regions of range overlap, we hypothesize that their songs will be more distinguishable than those of subspecies pairs that do not overlap. With greater song distinction, individuals in regions of overlap that encounter other subspecies may be able to more accurately identify members of their own subspecies from those of other subspecies.

A 2014 study suggests that dark-eyed juncos are able to discriminate their subspecies songs from those of other subspecies: male dark-eyed juncos in Virginia responded more strongly to playbacks of locally recorded songs than to songs recorded in California of a different subspecies [24]. Song discrimination between subspecies has also been demonstrated in a subset of other songbirds [25]; for example, male swamp sparrows respond more strongly to playbacks of songs from their subspecies than those from other subspecies, suggesting the possibility of continued divergence through song [26]. In contrast, subspecies of other songbird species have not shown song discrimination capability; for instance, a subspecies of the gray-breasted wood wren responds with similar strength to playbacks of its own song and the song of another subspecies [27].

Here, we explore song variation across the six majorly recognized subspecies of *Junco hyemalis*: *hyemalis, oreganus, caniceps, dorsalis, aikeni*, and *mearnsi*. Using song recordings collected from community-science databases, we parsed and analyzed dark-eyed junco songs using Chipper, a semi-automated song segmentation and analysis software [28]. Then, we compared the song features that were extracted from each song to assess whether subspecies exhibit significant song differences. In addition, we built a machine learning classifier trained on song data from across the six subspecies to determine whether subspecies’ songs can be readily distinguished. We also used song distinguishability metrics to determine whether the songs of subspecies pairs that are known to hybridize are more or less distinct than songs of subspecies pairs not known to hybridize. Finally, we measured pairwise genetic differences between the subspecies and assess whether the degree of genetic differentiation is associated with song diversification, geographic distance, or both.

## METHODS

### Attaining songs for analysis

We acquired dark-eyed junco song recordings from Macaulay Library [29] by requesting all available song files, as well as from Xeno-canto [30] using the WarbleR package (version 1.1.26) [31] in R. Macaulay Library and Xeno-canto are digital repositories of media captured from nature, including birdsong audio recordings. For our analyses, we compiled metadata for each recording, including subspecies, latitude, longitude, date, and time. Often, recordists include this relevant information when uploading song recordings. For recordings that listed a location but no geographic coordinates, we approximated the latitude and longitude by searching the location in Google Maps.

After obtaining the song recordings, we used version 3.1.3 of Audacity to manually extract individual song bouts (**Fig. 2A**). Most recordings contained multiple bouts, so we selected up to three bouts from each recording, preferentially choosing bouts that maximized signal-to-noise ratio and minimized vocalizations of other birds in the background. If all song bouts were equally clear, we extracted the first three. Some recordings yielded only one or two bouts due to short recording duration or poor audio quality. On rare occasions, we were unable to identify any suitable song bouts, which resulted in omission of the recording from the study.

**Figure 2.**
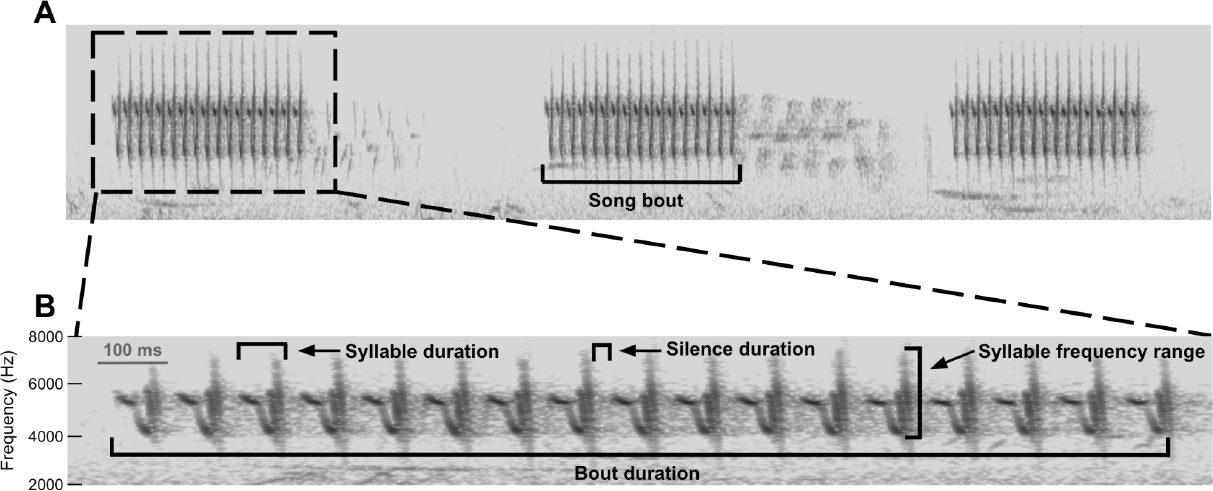
(A) Dark-eyed junco recording with multiple song bouts. (B) Spectrogram of an extracted song bout. Each song can be segmented into syllables and is characterized by several song features, such as bout duration, average syllable and silence duration, and average syllable frequency range.

Dark-eyed juncos have been observed singing two main song types: long-range songs and short-range songs [32]. The overwhelming majority of compiled recordings were long-range songs, so we exclusively studied this song type.

In total, we obtained 1,658 song recordings from Macaulay Library and Xeno-canto, and we were able to extract at least one song bout from 1,318 of these recordings, with 1,999 song bouts retained for analysis (**Table 1**). (A full list of song recordings and corresponding metadata is in **Supplemental Data 1**.)

**Table 1.**
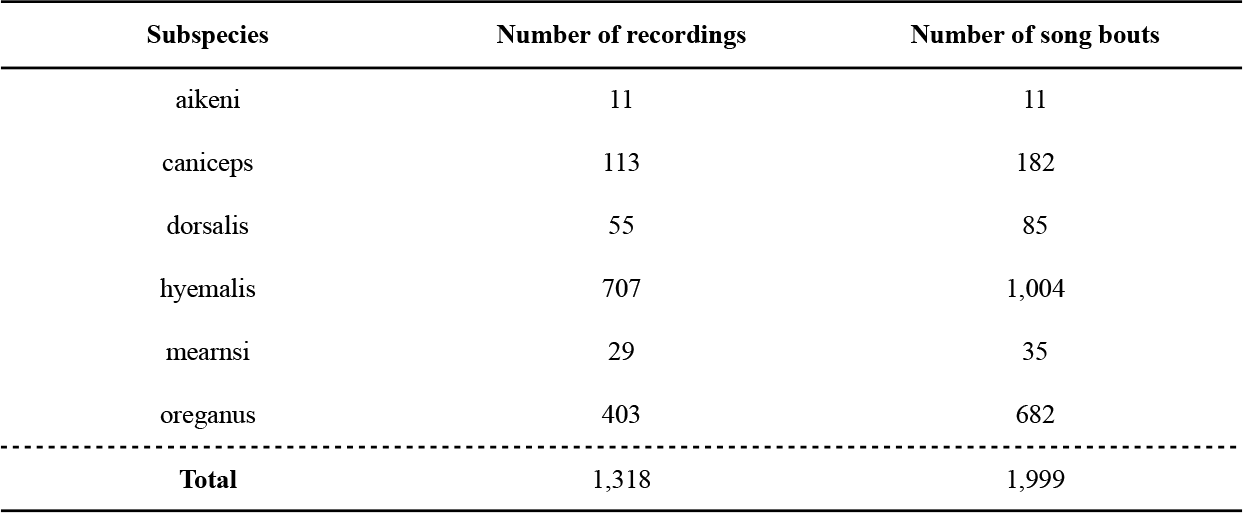
The number of recordings and song bouts obtained for each subspecies.

| Subspecies | Number of recordings | Number of song bouts |
| --- | --- | --- |
| aikeni | 11 | 11 |
| caniceps | 113 | 182 |
| dorsalis | 55 | 85 |
| hyemalis | 707 | 1,004 |
| mearnsi | 29 | 35 |
| oreganus | 403 | 682 |
| **Total** | 1,318 | 1,999 |

### Song segmentation and quantification of song features

To parse and analyze songs, we used Chipper [28], a graphical user interface that first allows users to supervise an automated syllable segmentation process and then analyzes properties of those syllables. In Chipper, songs are displayed as spectrograms—a visual representation showing frequencies of a sound signal over time (**Fig. 2B**). Chipper parses songs into syllables by identifying periods of sound separated by silences. To prevent lower amplitude syllables from being segmented incorrectly, we normalized amplitude across each song. To improve segmentation, we adjusted the signal-to-noise threshold, minimum syllable duration, and minimum silence duration. If any syllables were incorrectly parsed after adjusting these settings, we manually modified the syllable segmentation. If a song could not be cleanly segmented due to poor audio quality or excessive background noise, the song was excluded from further analysis. We provide song recording metadata, song-feature data, and our codes for statistical analysis and data visualization at https://github.com/CreanzaLab/JuncoAnalysis.

After song segmentation, we examined a random subset of 47 songs to estimate the noise threshold, defined as the minimum size a cluster of sound must be in order to not be discarded as noise. The estimated noise threshold was 81, meaning clusters of noise that are at least 81 pixels on the spectrogram are retained as signal. Additionally, we used the same subset of songs to estimate the syllable similarity threshold, defined as the percent syllable overlap that determines whether two syllables are considered the same or different. The estimated syllable similarity threshold was 53.5, meaning syllables with at least 53.5% signal overlap are considered to be repetitions of the same syllable. These average threshold values were then used for analysis of all songs.

We used the song analysis function in Chipper, which uses the syllable segmentation information and the noise and syllable similarity thresholds to generate quantitative data for several song features. We used 10 of the song-feature outputs from Chipper: average syllable duration (ms), bout duration (ms), number of syllables, rate of syllable production (1/ms, measured as number of syllables divided by bout duration), average silence duration (ms), number of unique syllables, mean syllable stereotypy, average syllable upper frequency (Hz), average syllable lower frequency (Hz), and average syllable frequency range (Hz).

### Analysis of quantitative song-feature data

Using the song-feature data, we performed statistical analyses to determine if there are significant song-feature differences between pairs of subspecies. First, we determined that none of the song features were normally distributed, so we used the natural log of each feature going forward. For each of the 15 possible subspecies pairs, we performed Wilcoxon rank-sum tests for each of the ten song features to test for significant subspecies-level song differences. We used the Bonferroni correction for multiple comparisons to adjust the *P*-value of each test result with a threshold for significance of α = 0.05, which reduces the likelihood of false positive results.

To reduce the dimensionality of our song-feature data, we used principal component analysis (PCA) and linear discriminant analysis (LDA). Respectively, these analyses allowed us to assess the degree of subspecies-level song differences by visualizing whether the subspecies form clusters in principal component (PC) space and whether any pair of song features have the ability to discriminate song data into subspecies.

### Machine learning classifier

We applied machine learning to assess how accurately dark-eyed junco subspecies can be distinguished by their songs. We built a random forest classifier using the tidymodels meta-package in R [33]. The classifier is trained on 10 song features and uses 100 decision trees to make predictions of subspecies identity. We tried increasing the number of decision trees and found that more trees very slightly decreased prediction accuracy, possibly from overfitting (**Table S2**). For each round of classification, 75% of randomly selected songs were part of the training set and the remaining 25% of songs belonged to the test set. We downsampled the training set using the themis R package [34] to obtain a balanced sample of songs from each subspecies. We tested the classifier’s prediction accuracy for discriminating between all six subspecies and its prediction accuracy for discriminating between pairs of subspecies. For each round of classification, we constructed a variable importance plot using the vip R package [35] to determine which song features were most valuable for the classification predictions.

### Hybridizing subspecies song analysis

We downloaded eBird sightings for each dark-eyed junco subspecies and for all hybrids. Hybrid sightings are generally denoted in eBird as a cross between two subspecies (e.g. “Dark-eyed Junco (Oregon x Pink-sided) - *Junco hyemalis oreganus* x *mearnsi*”), but hybrids between slate-colored and Oregon juncos are often denoted as a separate subspecies, *Junco hyemalis cismontanus*, also known as the Cassiar junco. We restricted these sightings to those recorded between March and August, which is the general breeding season of the dark-eyed junco, and used the latitude and longitude coordinates provided with each eBird sighting to visualize the breeding-season sightings on a map (**Fig. 1**). Based on this sighting data, we sorted the 15 subspecies pairs into one of three groups: subspecies pairs that are known to hybridize, subspecies pairs whose breeding ranges are adjacent but are not known to hybridize, and subspecies pairs whose breeding ranges are not adjacent. (Without nearby breeding ranges, subspecies are unlikely to have the opportunity to hybridize regardless of whether they can distinguish their own subspecies.) We investigated whether song distinguishability differs between the three groups using the machine learning classifier’s discrimination accuracy and the area of overlap in PC space after a principal component analysis of the song-feature data.

### Genetic analysis

We studied the genes sequenced by Friis *et al. 2016* [36], which includes four regions of mtDNA: 343 base pairs (bp) of the hypervariable region 1 of the control region (CR); 640 bp of the cytochrome *c* oxidase subunit I (COI); 945 bp of the NADH dehydrogenase subunit 2 (ND2); and 870 bp of the ATPase genes 6 and 8 (ATPase). They also amplified 489 bp from intron 5 of the nuclear fibrinogen beta chain (FGB) gene. For each of these genes, we added additional sequences when available on NCBI. Additionally, we concatenated the ND2, COI, and ATPase genes from the same individuals for broader genetic analysis. The total number of sequences compiled for each gene is in **Table 2**. We used NCBI’s BatchEntrez tool to retrieve fasta files of the desired sequences, then aligned the sequences of each gene using MAFFT [37].

**Table 2.** The number of genetic sequences available per subspecies for each gene and the concatenated genetic data.

| Gene | Aikeni | Caniceps | Dorsalis | Hyemalis | Mearnsi | Oreganus | Total |
| --- | --- | --- | --- | --- | --- | --- | --- |
| CR | 8 | 9 | 10 | 7 | 8 | 36 | 78 |
| COI | 8 | 12 | 10 | 34 | 8 | 50 | 122 |
| ND2 | 8 | 9 | 10 | 6 | 8 | 36 | 77 |
| ATPase | 8 | 9 | 10 | 6 | 8 | 37 | 78 |
| FGB | 6 | 4 | 4 | 16 | 3 | 18 | 51 |
| Concatenated | 8 | 9 | 10 | 6 | 8 | 36 | 77 |

To analyze the genetic sequences, we used the software Arlequin (version 3.5.2.2) [38] to calculate pairwise fixation indices (F_ST_) between subspecies for each gene and the concatenated data. The thresholds for significance (α) were adjusted according to the Benjamini-Hochberg procedure, which limits false discovery rate [39]. Pairwise F_ST_ calculations quantify genetic differentiation between the subspecies by measuring the proportion of genetic variation that is due to allelic differences within versus between populations. Using Arlequin, we also performed Analyses of Molecular Variance (AMOVA) for each gene and the concatenated data to quantify the amount of genetic variation that is due to differences in molecular markers within and between populations. Finally, we used the adegenet R package [40] to run principal component analyses (PCA) for each of the five genes and the concatenated data to assess whether the genetic data were clustered by subspecies in PC space, which would suggest genetic differentiation between the subspecies.

### Joint analysis between song, genetic, and geographic data

To elucidate significant relationships between our song, genetic, and geographic data, we used Procrustes analyses to measure the spatial similarity between each pair of datasets: 1.) song and geographic data, 2.) genetic and geographic data, and 3.) song and genetic data. For the song and genetic datasets, we used PC1 and PC2 from their respective principal component analyses. The geographic dataset consisted of longitude and latitude corresponding to the locations of individuals from the song and genetic data. Procrustes analyses rotate and transform a matrix to best superimpose it onto an equally-sized target matrix, such that the sum of squared-distances between the corresponding points of the transformed and target matrix are minimized. For each comparison, we used the ‘protest’ function in the vegan R package [41], which executes permutation tests by randomly permuting the order of one dataset and repeating the Procrustes to determine the significance of the original result.

## RESULTS

### Analysis of quantitative song-feature data

We tested whether song features differed significantly in their distribution between pairs of subspecies. The pairwise Wilcoxon rank-sum tests showed significant differences (P < α = 0.05) between at least one pair of subspecies for all song features except bout duration, number of unique syllables, and mean syllable stereotypy (**Fig. S1-S10**).

The principal component analysis (PCA) of the song-feature data revealed considerable overlap in PC space. Though the subspecies do not form distinct clusters, some subspecies are more localized than others in PC space, such as dorsalis and aikeni. PC1 and PC2 respectively explain 33.4% and 19.11% of the variance in the data (**Fig. 5A**). The linear discriminant analysis (LDA) showed similar patterns, with no distinct subspecies clusters (**Fig. S11**). The average silence duration and mean syllable stereotypy were the song features with the greatest ability to separate the subspecies, with a moderate discriminative power of 0.3345. The lack of distinct subspecies clusters from the PCA and LDA suggests an absence of subspecies-level song diversification.

### Machine learning classifier

The random forest classifier had an average accuracy of 75.8% when distinguishing between pairs of subspecies (**Table S3**). When distinguishing between all six subspecies, the classifier had an average accuracy of 28.9%. The sizes of the training sets varied between subspecies pairs because of downsampling. However, the classifier’s performance is not correlated with training set size (**Fig. S12**). The variable importance plots showed that the most important song features the classifier used to categorize test songs into subspecies were rate of syllable production and average syllable lower frequency (**Table S4**).

### Hybridizing subspecies song analysis

Hybrid sightings are rare, accounting for less than 1% of total sightings in regions of overlap between the subspecies (**Table S1**). Subspecies pairs with observed hybrids are *hyemalis* x *oreganus* (n = 95 hybrid sightings), *caniceps* x *dorsalis* (n = 5), *mearnsi* x *caniceps* (n = 61), *oreganus* x *caniceps* (n = 7), *mearnsi* x *aikeni* (n = 5), and *oreganus* x *mearnsi* (n = 6). Subspecies pairs with adjacent ranges that do not appear to hybridize are *oreganus* x *aikeni, caniceps* x *aikeni*, and *oreganus* x *dorsalis*. Subspecies pairs that are known to hybridize have an average classifier accuracy of 69.6%, pairs with non-adjacent ranges have a moderate average classifier accuracy of 75.0%, and pairs with adjacent ranges that are not known to hybridize have the greatest average classifier accuracy, 90.0% (**Fig. 3A**), indicating that subspecies pairs that avoid hybridizing also have more distinguishable songs. We also quantified the proportion of song-feature PCA overlap, which gives an indication of the amount of shared song variation between the subspecies. With this measure, subspecies pairs that are known to hybridize have an average PCA overlap of 46.7%, pairs with non-adjacent ranges have an average PCA overlap of 30.3%, and pairs with adjacent ranges that are not known to hybridize have an average PCA overlap of 32.0%, suggesting that subspecies pairs that avoid hybridizing also have less overlapping song variation.

**Figure 3.**
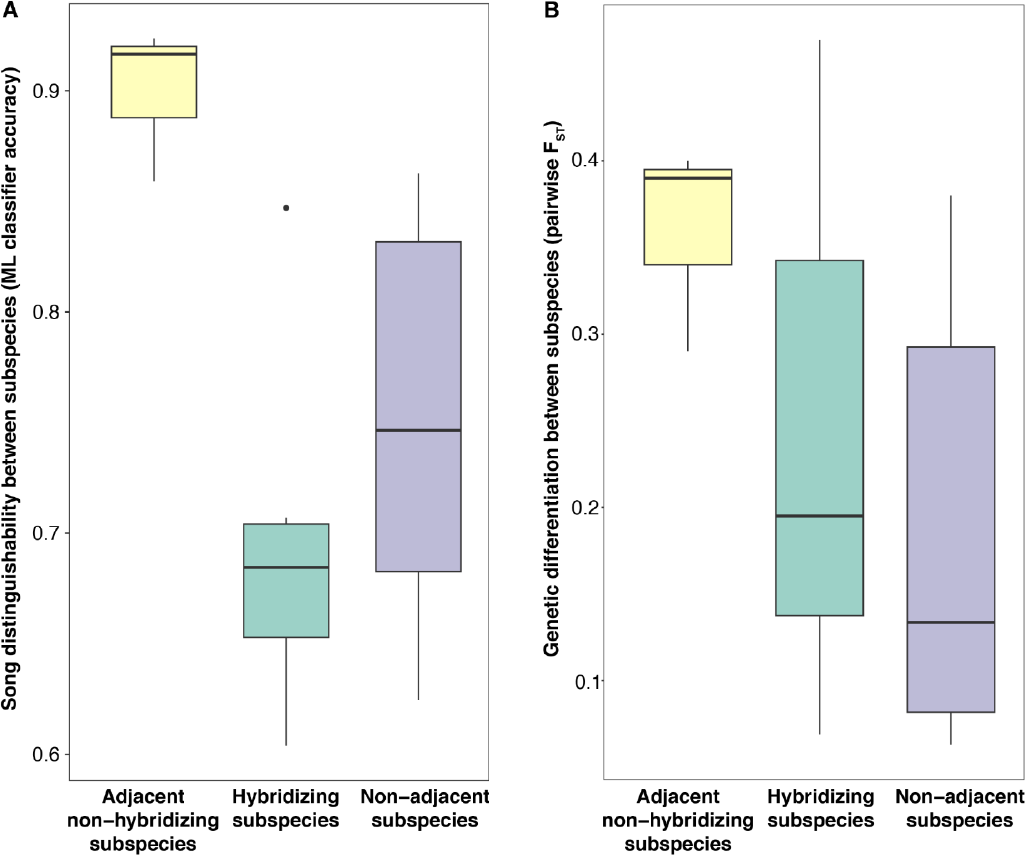
Results of machine learning song-feature analysis and genetic analysis, partitioned by whether subspecies appear to hybridize at the boundaries of their adjacent ranges. The partitioned groups are subspecies pairs that have adjacent ranges and are not known to hybridize (n = 3), pairs that are known to hybridize (n = 6), and pairs that do not have adjacent ranges (n = 6). (A) Distribution of average machine learning classifier accuracy for the partitioned groups. (B) Distribution of pairwise F_ST_ for the partitioned groups.

### Genetic analysis

The pairwise F_ST_ calculations show that the concatenated genetic data and all genes except ND2 have at least one pair of subspecies with a significant F_ST_ (**Fig. 4, Figs. S13-S17**). We found that pairwise F_ST_ was greatest for subspecies pairs that have an adjacent range but do not hybridize, indicating greater genetic differentiation than we observed in pairs that are known to hybridize and pairs that do not share an adjacent range (**Fig. 3B**). The AMOVA tests reveal the amount of genetic variance between and within subspecies using molecular markers. For the concatenated genetic data, 27.3% of the variance is between subspecies and 72.7% of the variance is within subspecies. (**Table 3, Tables S5-S9**). The subspecies assignments of the genetic data were permuted and an empirical *P*-value of *P* <1.0 × 10^−4^ was calculated as the probability that the F_ST_ of the permuted data is greater than the F_ST_ of the observed data. The PCAs of the genetic data suggest some genetic population structure, but do not show fully separated subspecies clusters (**Fig. 6A, Figs. S18-22**).

**Table 3.**
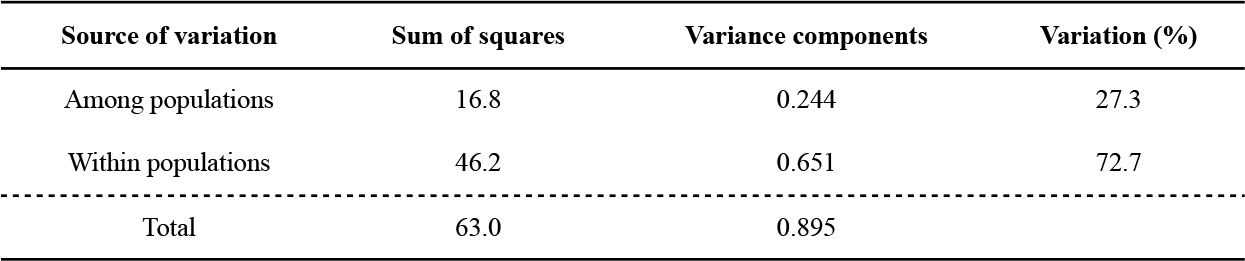
AMOVA results for the concatenated genetic data. 27.3% of the variance was between subspecies and 72.7% of the variance was within subspecies. The genetic data were permuted and an empirical *P*-value of *P* < 1.0 × 10^−4^ was calculated as the probability that the F_ST_ of the permuted data is greater than the F_ST_ of the observed data.

| Source of variation | Sum of squares | Variance components | Variation (%) |
| --- | --- | --- | --- |
| Among populations | 16.8 | 0.244 | 27.3 |
| Within populations | 46.2 | 0.651 | 72.7 |
| Total | 63.0 | 0.895 |  |

**Figure 4.**
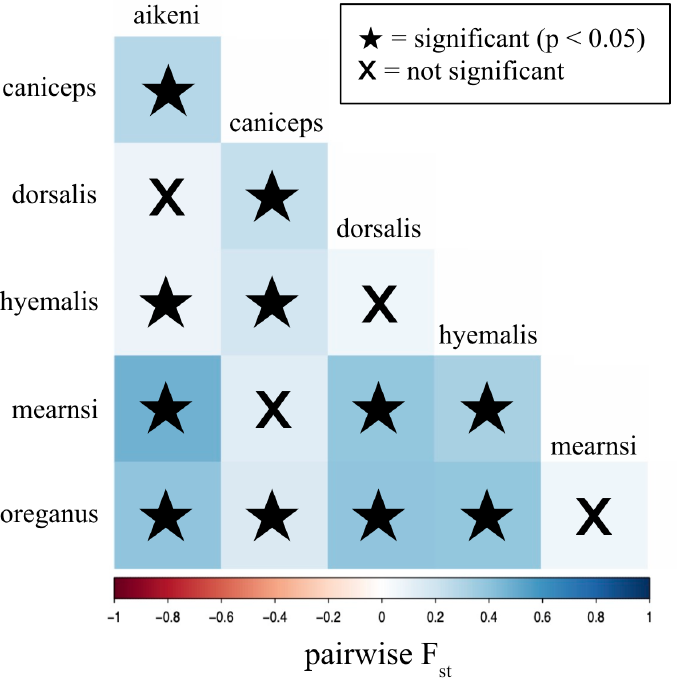
Pairwise F_ST_ matrix for the concatenated genetic data. Subspecies pairs with significant F_ST_ are denoted with stars. The color scale corresponds to F_ST_, and thus, the relative degree of genetic differentiation.

**Figure 5.**
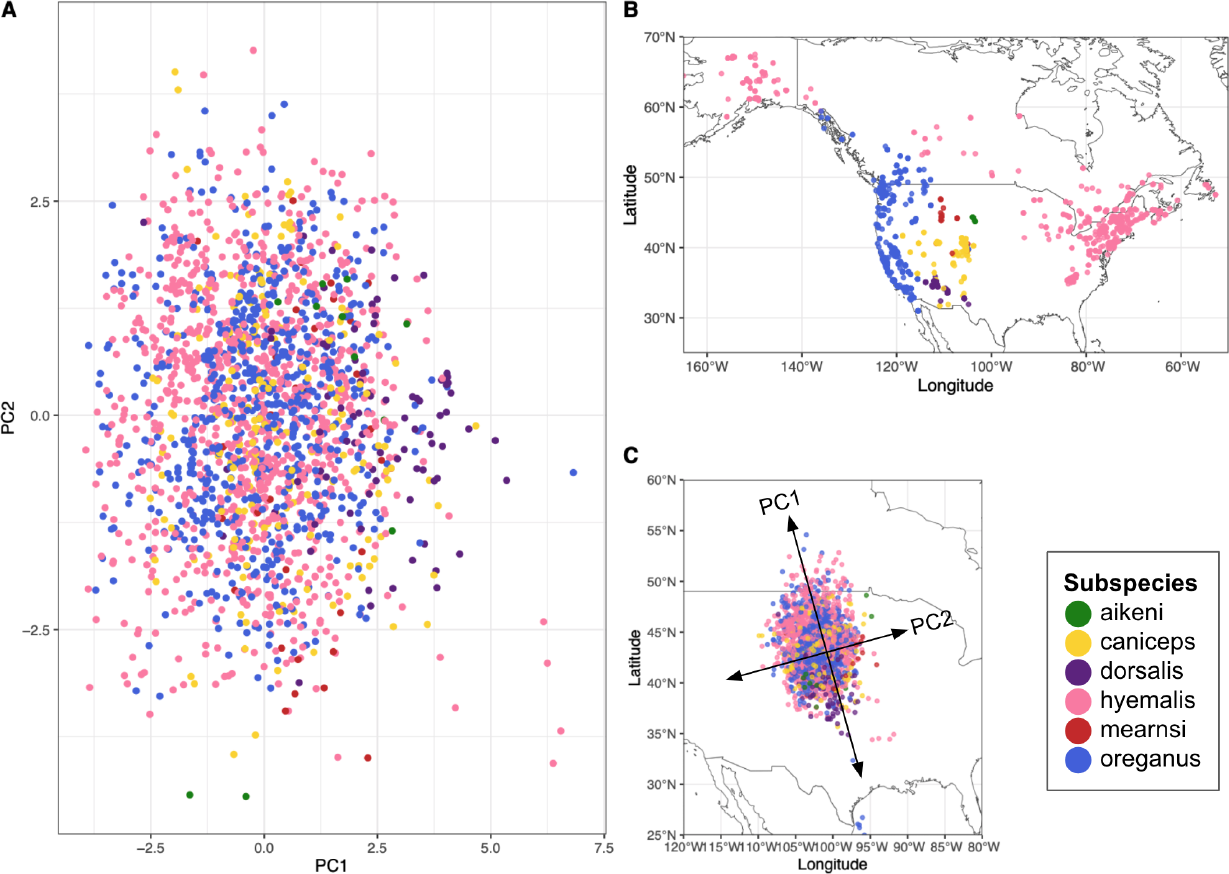
Spatial distribution of song data. (A) PC1 and PC2 of the song data using 10 song features, colored by subspecies. (B) Map displaying the locations of song recordings. (C) Procrustes analysis comparing PC1 and PC2 of the song data to longitude-latitude coordinates. Procrustes rotation = 0.155.

**Figure 6.**
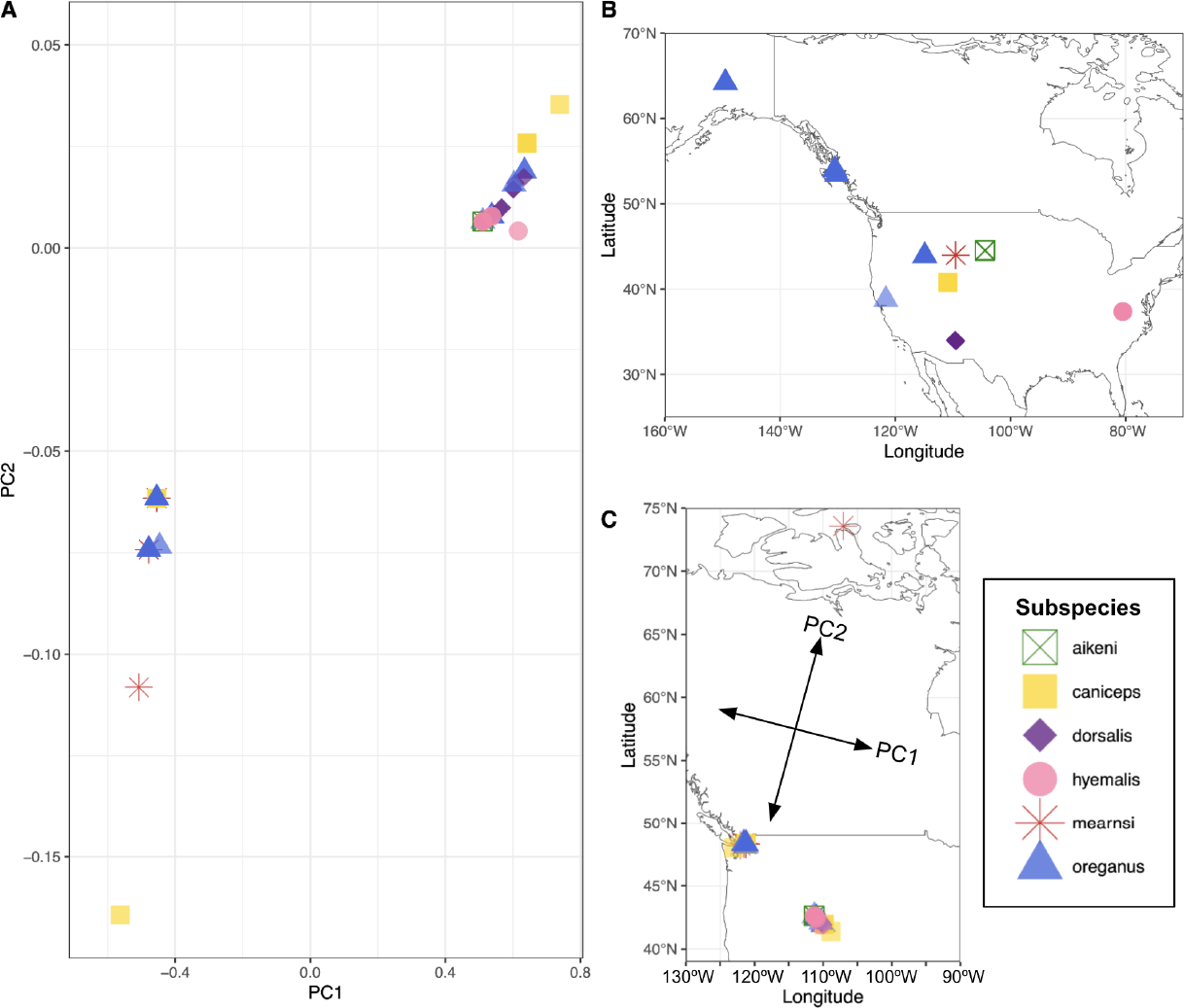
Spatial distribution of the concatenated genetic data. (A) PC1 and PC2 of the concatenated genetic data, colored by subspecies. (B) Map displaying the locations of where genetic samples were taken. (C) Procrustes analysis comparing PC1 and PC2 of the genetic data to longitude-latitude coordinates. Procrustes rotation = 0.356.

### Joint analysis between song, genetic, and geographic data

With Procrustes analyses, we constructed a two-dimensional plot of each data type—PC1 vs. PC2 of the song data, PC1 vs. PC2 of the genetic data, and latitude vs. longitude—and compared these plots to assess whether each data pair showed a similar spatial structure. We found a significant spatial relationship between song and geographic data (*P* = 1.0 × 10^−3^) and between genetic and geographic data (*P* = 6.0 × 10^−4^) but not between song and genetic data (*P* = 0.292).

## DISCUSSION

The dark-eyed junco has been a well-studied species for nearly a century, beginning with migration experiments in 1924 [42]. Though the dark-eyed junco subspecies have phenotypic differentiation in plumage and morphology, they are known to interbreed in areas of overlap, and they are generally considered one species [16]. However, since these subspecies re-colonized much of North America after the Last Glacial Maximum (∼18,000 years ago), their phenotypic and genetic differences represent an extremely rapid case of diversification [36]. Visual plumage and morphological differences as well as geographic distribution likely contribute to mating occurring disproportionately within subspecies, but song could also play a role in maintaining subspecies boundaries, as song is used in species recognition and mate attraction. Culturally transmitted traits like song are important to study because they can evolve faster than genetically heritable traits and accelerate evolution and speciation [14]. No previous studies have extensively assessed song differences between dark-eyed junco subspecies. This study aims to determine if there are significant song differences between dark-eyed junco subspecies and the role potential song differences may play in maintaining reproductive boundaries between subspecies.

Our pairwise comparisons of dark-eyed junco song features revealed that seven of the ten studied features have significant differences between at least one pair of subspecies. Number of unique syllables, mean syllable stereotypy, and bout duration do not have significant differences between subspecies pairs. The lack of significant differences in these song features makes sense; dark-eyed junco songs typically consists of a single repeated syllable, which results in low variability in these three song features between subspecies. For song features with significant differences, the number of subspecies pairs with significant differences varies. Song features with ≥8 out of 15 significantly different subspecies pairs include number of syllables, rate of syllable production, and average syllable lower frequency. These song features have more plasticity than other song features that are more constrained by the general characteristics of dark-eyed junco songs. Some subspecies pairs do not have any song features that significantly differ (*hyemalis*/*oreganus* and *aikeni*/*mearnsi*), while other pairs have as many as 6 song features with significant differences (*hyemalis*/*dorsalis, oreganus*/*aikeni*, and *oreganus*/*dorsalis*). Interestingly, subspecies pairs with no significantly different song features are known to hybridize while pairs with the greatest number of significantly different song features are not known to hybridize.

While significant differences exist in individual song features, collectively the differences are not substantial enough to consistently classify songs by subspecies. The PCA and LDA of the song data that considered all 10 studied song features were unable to clearly separate the subspecies into clusters graphically. Additionally, the random forest classifier was unable to unequivocally predict the subspecies of test songs. When discriminating between all six subspecies, the classifier was only able to predict the subspecies of test data with 28.9% accuracy. In nature, the likelihood of a dark-eyed junco having to distinguish between all six subspecies is unlikely; most areas of overlap have interaction between two, or rarely three, subspecies. The classifier was able to discriminate between pairs of subspecies with an average accuracy of 75.8%.

Interestingly, when we partitioned the subspecies pairs based on whether or not they hybridized, we found a clear pattern: subspecies pairs in adjacent ranges that did not hybridize had much more distinct songs than subspecies pairs that did hybridize. The proportion of song-feature PCA overlap, the random forest classifier prediction accuracies, and the degree of song-feature differences all support this finding. The average proportion of PCA overlap is 46.7% for subspecies pairs that hybridize and 32.0% for pairs that have adjacent ranges but do not hybridize (Table **S10**). A greater proportion of PCA overlap indicates that the subspecies songs are less distinct when considering the variation in all 10 studied song features. Similarly, the random forest model was able to discriminate between songs of hybridizing pairs with 69.6% accuracy but discriminated between songs of pairs that have adjacent ranges that do not hybridize with 90.0% accuracy. These results indicate that songs of hybridizing subspecies are less distinguishable than songs of subspecies pairs that have adjacent ranges but do not hybridize. Thus, song distinguishability appears to have predictive power for determining which subspecies will hybridize, suggesting that song might play a role in reinforcing subspecies boundaries. In the same vein, lower song distinguishability in the pairs that are known to hybridize may allow a male of one subspecies to occasionally attract a female of another subspecies, leading to hybridization. In summary, song might play a role in reinforcing certain subspecies boundaries more than others.

The significant pairwise F_ST_ values support the previous finding of genetic differentiation between dark-eyed junco subspecies [36]. Pairwise F_ST_ values were greater for subspecies pairs that have adjacent ranges and are not known to hybridize, suggesting that these pairs are more reproductively isolated than subspecies pairs known to hybridize.

From the Procrustes analyses, we found geographic signal in both the song and genetic data. Thus, individuals who share a range are more likely to share song characteristics and more likely to be genetically similar. Further, geographic signal suggests that song and genetic changes may accumulate in a neutral manner with geographic distance, such that plumage and morphology might be sufficient for local populations to distinguish one another visually.

Though this research suggests that song differences might reinforce certain subspecies boundaries more than others, there may be other explanations for the song variation observed within and between subspecies. For example, the environment has been shown to influence the evolution of mating signals like birdsong through selection acting directly on signal transmission or through morphological adaptations to an environment (like body and beak size), which indirectly affect signal production [43]. Further research is needed to investigate whether environmental or morphological conditions can explain differences in dark-eyed junco songs. Additionally, playback experiments could more directly test how well individual dark-eyed juncos can distinguish between songs of their subspecies and songs of other subspecies. The random forest classifier trained on quantitative song-feature data is a proxy for song distinguishability, but playback experiments generally provide a stronger measure of subspecies recognition than acoustic trait analyses [25].

In sum, we analyze community-science recordings from across North America and show that there is geographic signal in both song and genetic variation in the dark-eyed junco and that there are significant song-feature differences in several pairs of subspecies. These song differences do not allow us to unequivocally distinguish all subspecies from one another by their songs. However, when we consider pairs of subspecies that are likely to interact in nature, we observe a consistent pattern: the songs of adjacent subspecies pairs with no observed hybrids have songs that are more distinguishable from one another than subspecies pairs that produce documented hybrids, and genetic differentiation is also higher in the adjacent pairs that do not hybridize. This pattern suggests that subspecies could use learned song differences as a reproductive isolation mechanism. For subspecies that do hybridize, between-subspecies hybrids account for less than 1% of documented junco sightings in their regions of overlap even though their songs are difficult to distinguish, suggesting that subspecies use other signals to distinguish themselves as well, such as plumage differences. Taken together, our results suggest both that reproductive isolation between subspecies can be a nuanced phenomenon and that community-science data can be analyzed to shed light on the speciation process.

## ACKNOWLEDGEMENTS

We are grateful for song recording media from The Macaulay Library at the Cornell Lab of Ornithology (www.macaulaylibrary.org) and Xeno-canto (www.xeno-canto.org). A full list of the song recordings used from these digital repositories and their metadata can be found at https://www.github.com/CreanzaLab/JuncoAnalysis.

## Notes

https://github.com/CreanzaLab/JuncoAnalysis

### Competing Interest Statement

The authors have declared no competing interest.

