## Supplemental Materials for "Song distinguishability predicts reproductive isolation between subspecies of the dark-eyed junco"

**Table S1** Number of hybrid sightings and the number of parent subspecies sightings in hybridizing subspecies pairs' regions of overlap. The hybrid-to-parent sightings ratio is the number of hybrid sightings divided by the number of parent sightings in the region of overlap.

| Hybridizing subspecies pair | Number of hybrid sightings | Subspecies 1 sightings in region of overlap | Subspecies 2 sightings in region of overlap | Hybrid:parent sightings ratio |
|---|---|---|---|---|
| hyemalis x oreganus | 95 | 2,079 | 13,645 | 0.00604 |
| mearnsi x caniceps | 61 | 1,656 | 1,653 | 0.01843 |
| mearnsi x aikeni | 5 | 191 | 26 | 0.02304 |
| caniceps x dorsalis | 5 | 291 | 435 | 0.00689 |
| oreganus x caniceps | 7 | 270 | 2,112 | 0.00294 |
| oreganus x mearnsi | 6 | 2,752 | 3,254 | 0.00100 |
| Average: | | | | 0.00972 |

**Table S2** Random forest classifier's discrimination between hyemalis and oreganus songs is largely unaffected by increases in the number of decision trees. Accuracy decreases very slightly with more decision trees, suggesting possible overfitting. Therefore, we used 100 decision trees for our random forest classifier.

| Number of decision trees | hyemalis accuracy | oreganus accuracy | Averaged accuracy |
|---|---|---|---|
| 100 | 0.705 | 0.612 | 65.9% |
| 150 | 0.705 | 0.594 | 65.0% |
| 200 | 0.709 | 0.594 | 65.2% |
| 500 | 0.713 | 0.571 | 64.2% |
| 1000 | 0.709 | 0.588 | 64.9% |

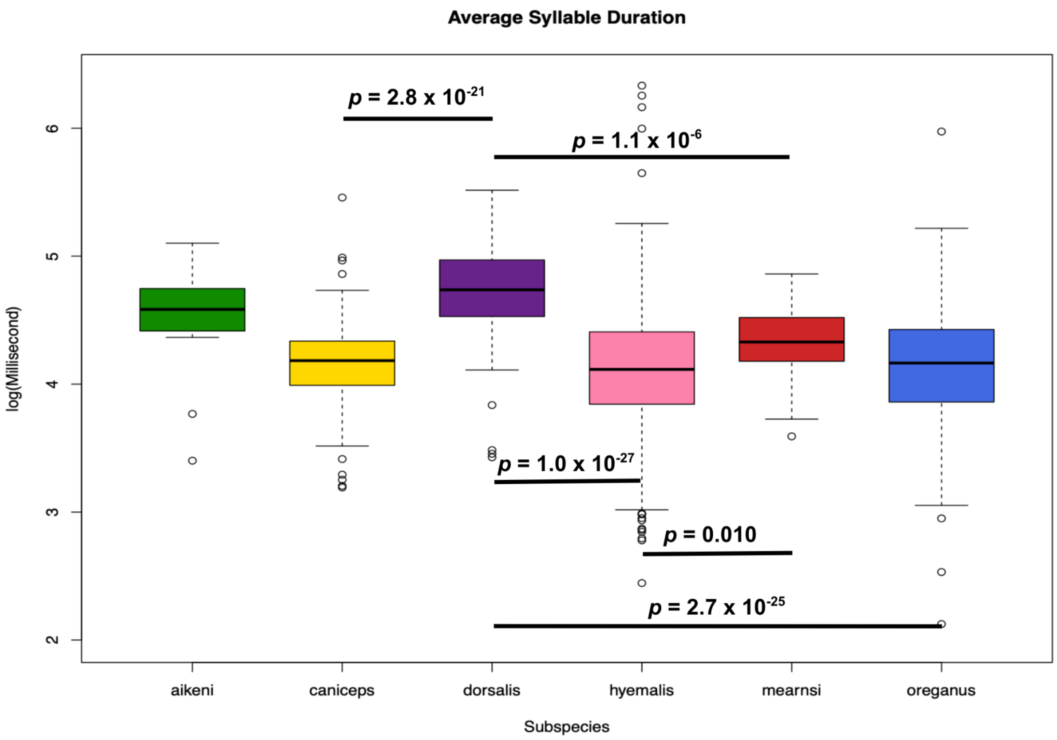


**Figure S1** Average syllable duration distribution for individuals across subspecies. Significantly different pairs for this song feature are denoted. *P*-values are Bonferroni-adjusted and $\alpha = 0.05$.

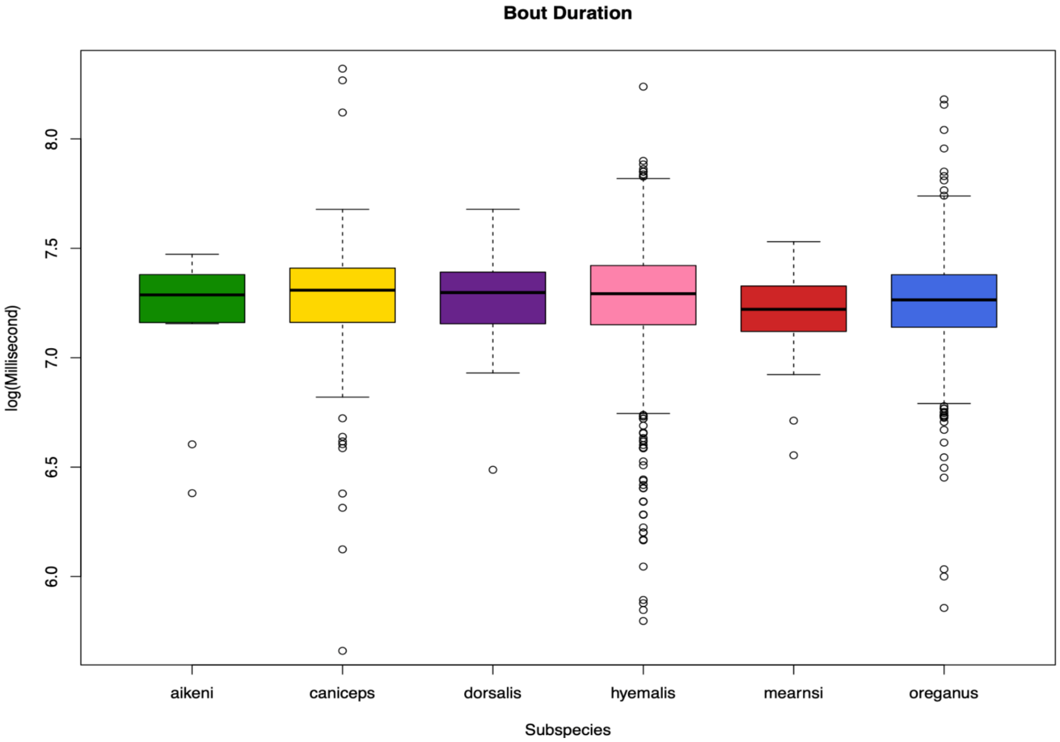


**Figure S2** Bout duration distribution for individuals across subspecies. There are no significantly different pairs for this song feature.

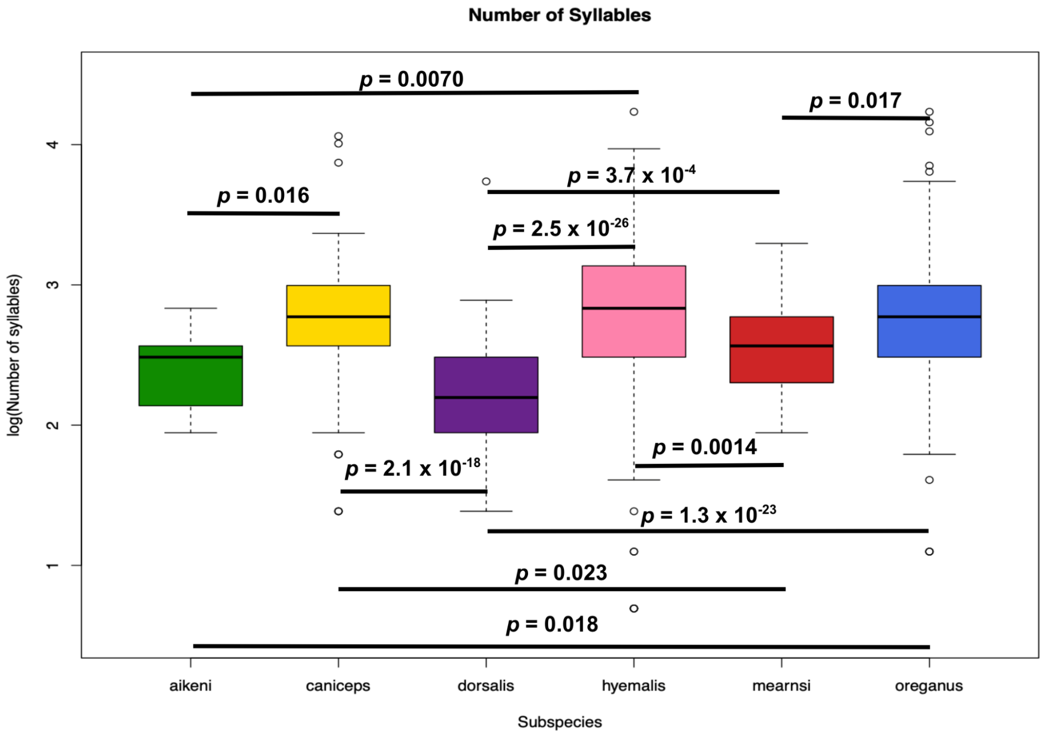


**Figure S3** Number of syllables distribution for individuals across subspecies. Significantly different pairs for this song feature are denoted. *P*-values are Bonferroni-adjusted and $\alpha = 0.05$.

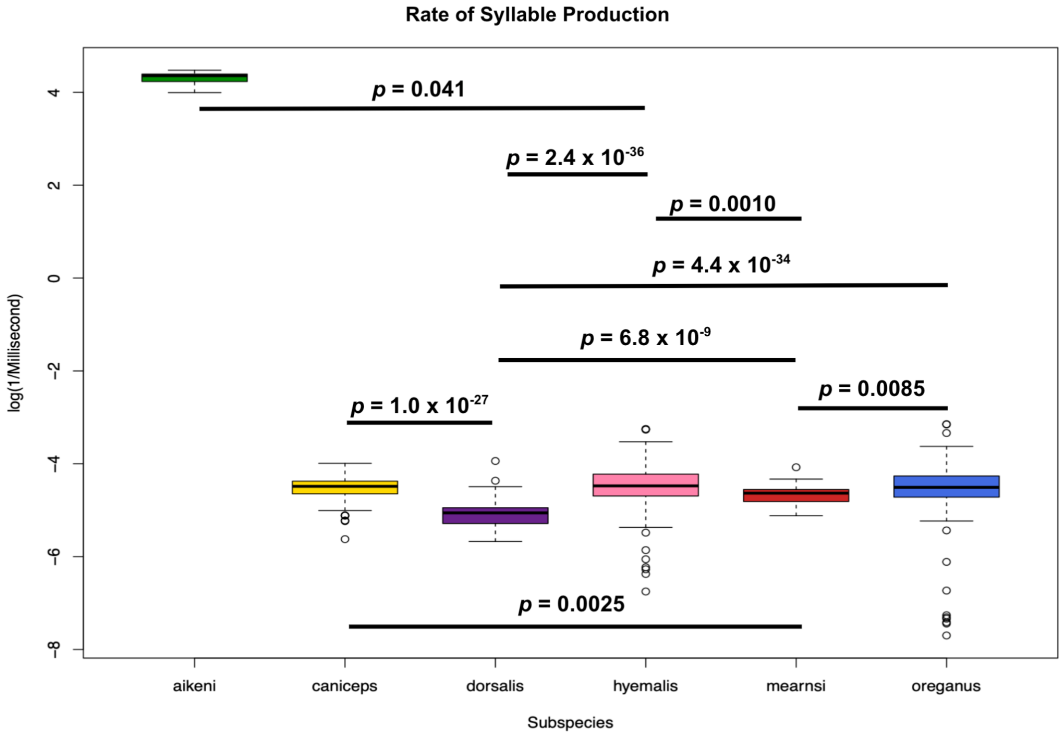


**Figure S4** Rate of syllable production for individuals across subspecies. Significantly different pairs for this song feature are denoted. *P*-values are Bonferroni-adjusted and $\alpha = 0.05$.

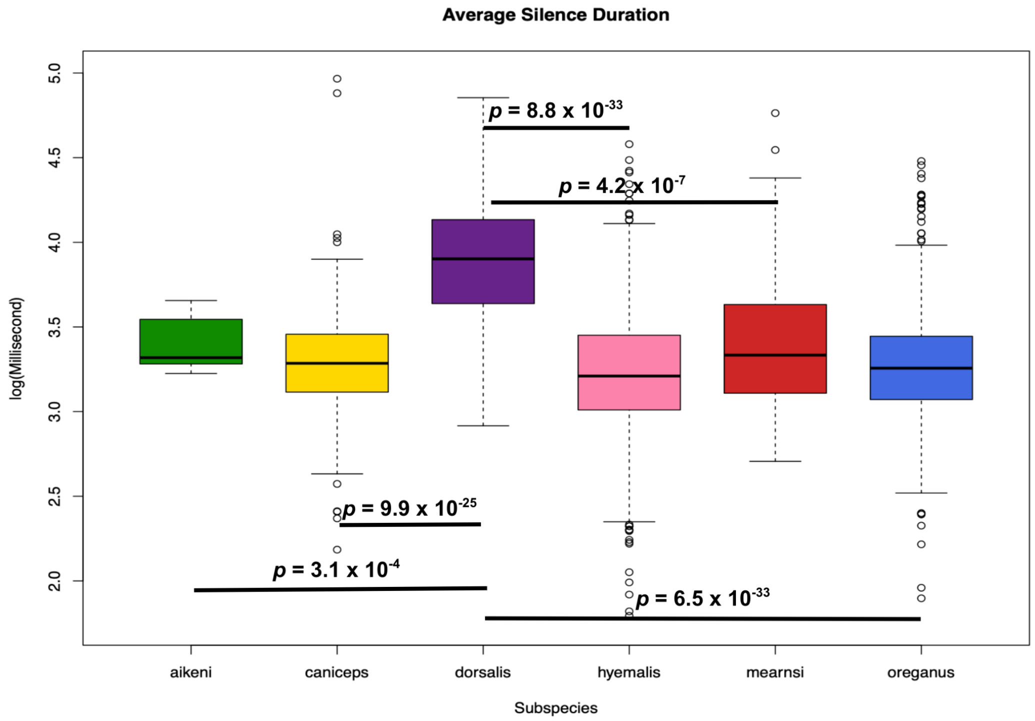


**Figure S5** Average silence duration distribution for individuals across subspecies. Significantly different pairs for this song feature are denoted. *P*-values are Bonferroni-adjusted and $\alpha = 0.05$.

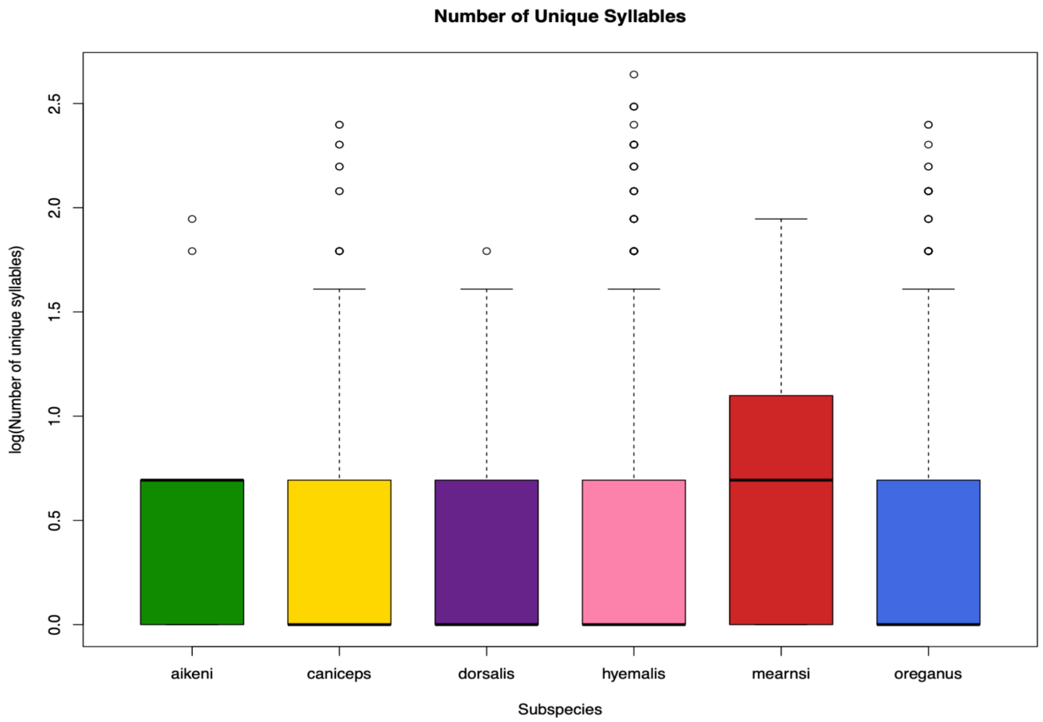


**Figure S6** Number of unique syllables distribution for individuals across subspecies. There are no significantly different pairs for this song feature.

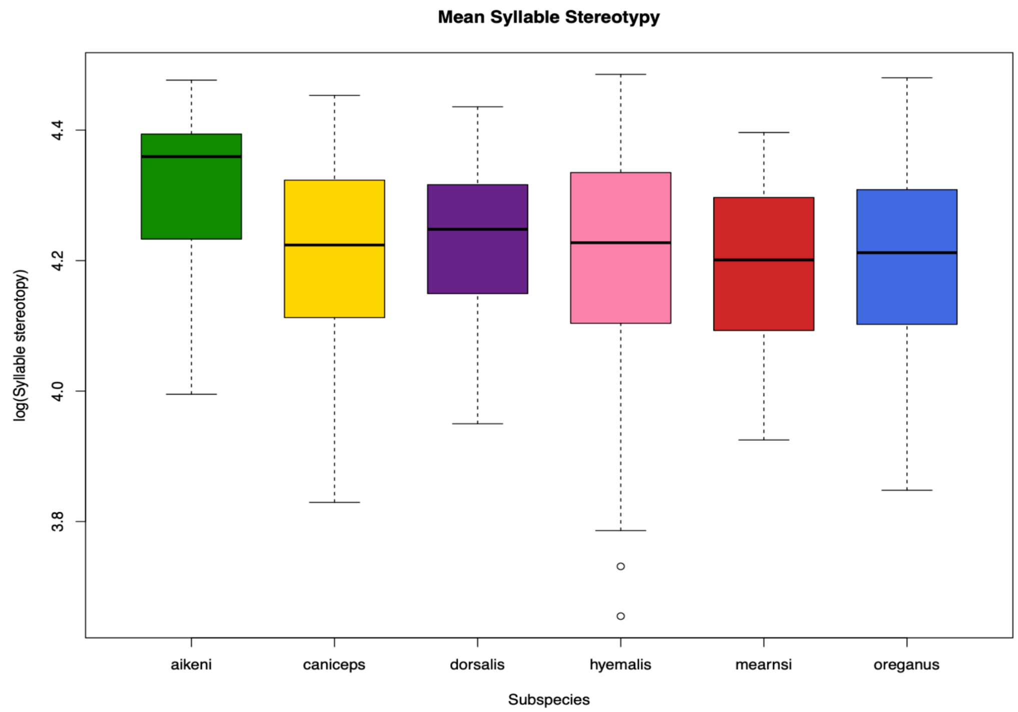


**Figure S7** Mean syllable stereotypy distribution for individuals across subspecies. There are no significantly different pairs for this song feature.

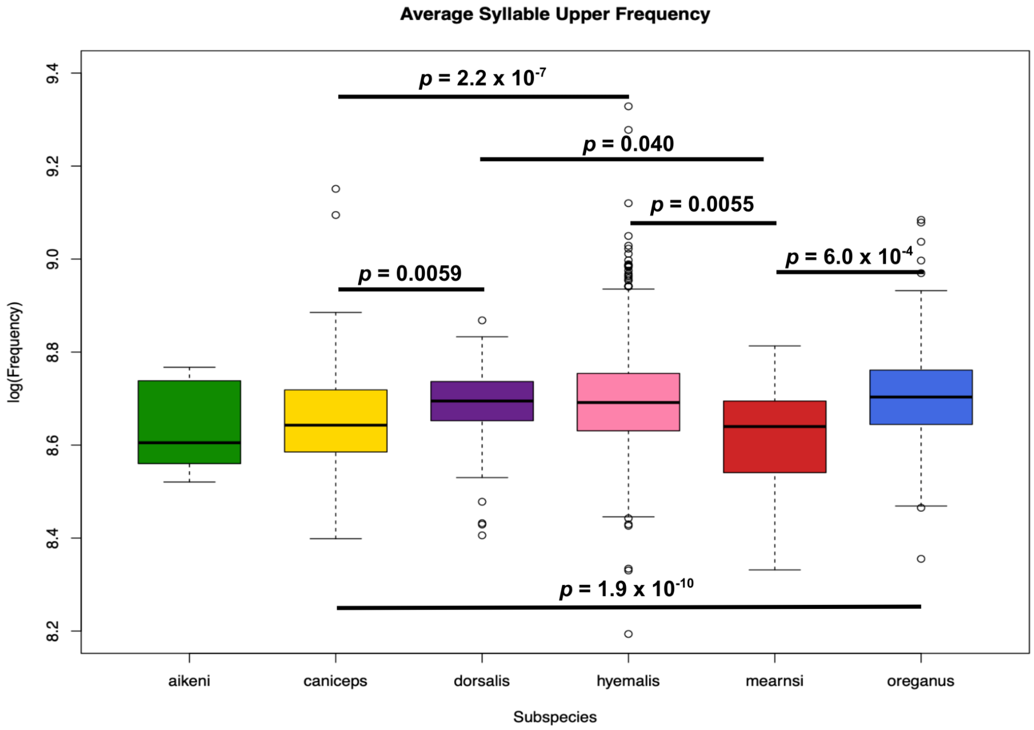


**Figure S8** Average syllable upper frequency distribution for individuals across subspecies. Significantly different pairs for this song feature are denoted. *P*-values are Bonferroni-adjusted and $\alpha = 0.05$.

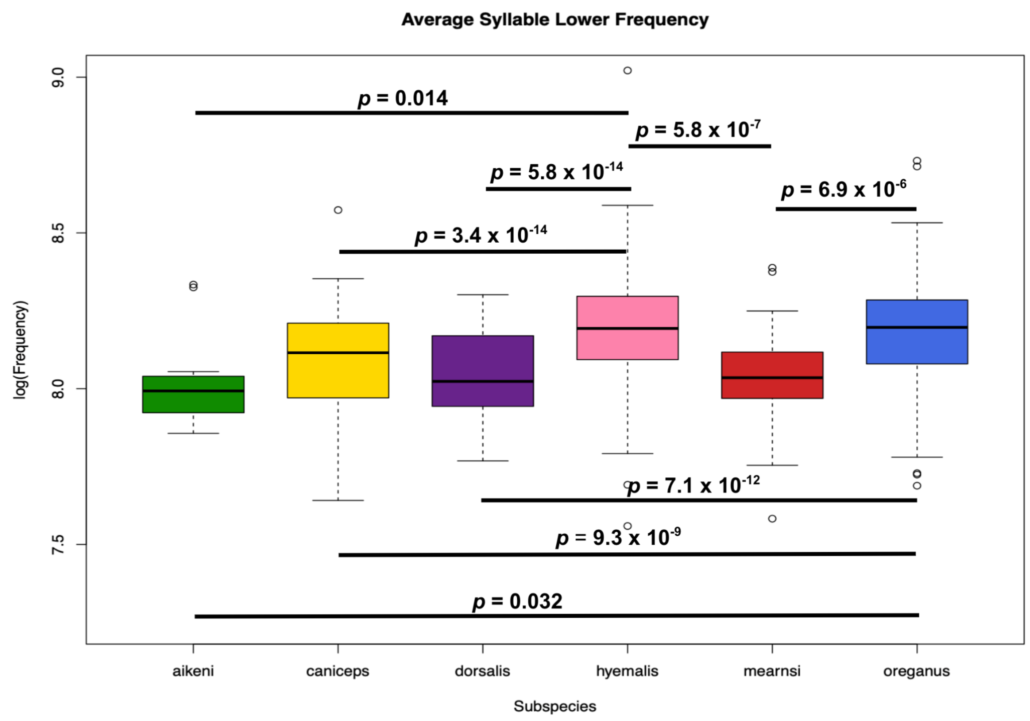


**Figure S9** Average syllable lower frequency distribution for individuals across subspecies. Significantly different pairs for this song feature are denoted. *P*-values are Bonferroni-adjusted and α = 0.05.

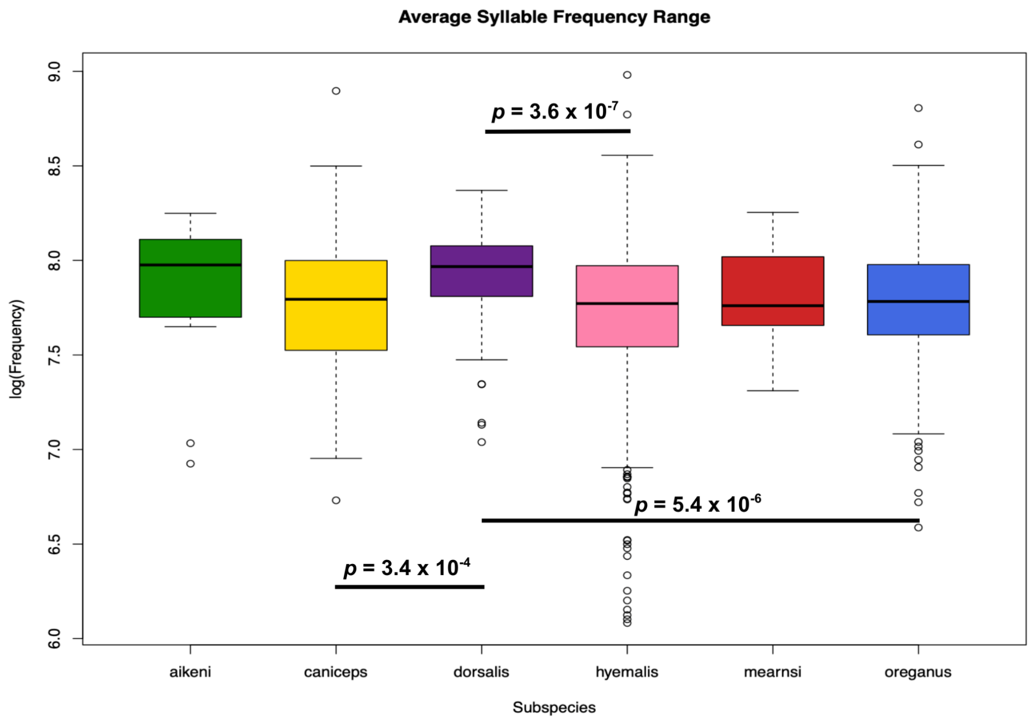


**Figure S10** Average syllable frequency range distribution for individuals across subspecies. Significantly different pairs for this song feature are denoted. *P*-values are Bonferroni-adjusted and α = 0.05.

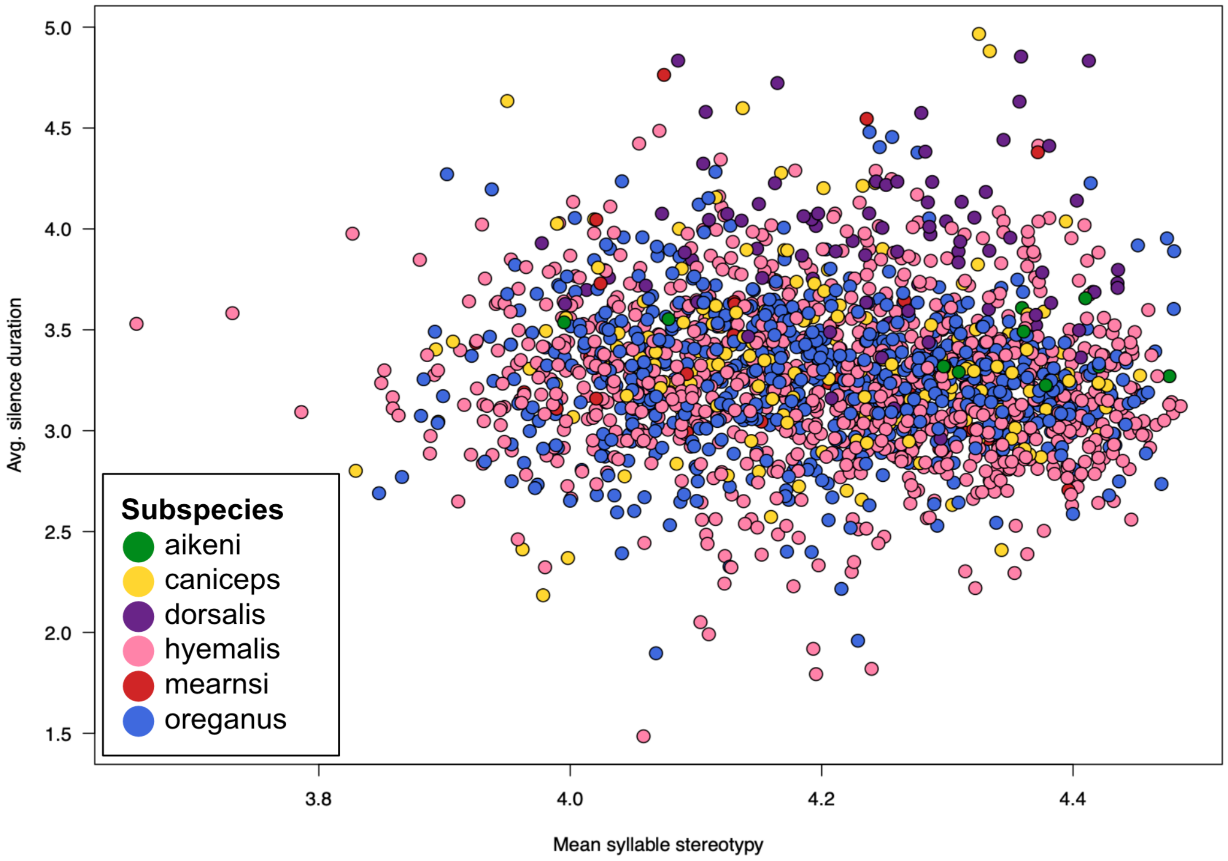


**Figure S11** Linear discriminant analysis (LDA) for the song data. After testing every pair of song features, the song feature pair that was able to best discriminate the data was mean syllable stereotypy and average syllable duration, with a discriminative power of 0.3345.

**Table S3** Random forest classifier's accuracies for distinguishing between pairs of subspecies.
Subspecies pair with adjacent ranges that are not known to hybridize
Subspecies pair with hybrid sightings
Subspecies pair without adjacent ranges that are not known to hybridize

| Subspecies pair | Training Set Size | Subspecies 1 Accuracy | Subspecies 2 Accuracy | Averaged accuracy |
|---|---|---|---|---|
| oreganus \| aikeni | 9 | 0.718 | 1.00 | 85.9% |
| caniceps \| aikeni | 10 | 0.833 | 1.00 | 91.7% |
| dorsalis \| oreganus | 59 | 0.900 | 0.947 | 92.4% |
| mearnsi \| oreganus | 25 | 0.571 | 0.637 | 60.4% |
| mearnsi \| aikeni | 8 | 0.625 | 0.667 | 64.6% |
| hyemalis \| oreganus | 507 | 0.659 | 0.688 | 67.4% |
| caniceps \| mearnsi | 24 | 0.766 | 0.625 | 69.6% |
| caniceps \| oreganus | 138 | 0.702 | 0.712 | 70.7% |
| caniceps \| dorsalis | 59 | 0.894 | 0.800 | 84.7% |
| hyemalis \| mearnsi | 26 | 0.582 | 0.667 | 62.5% |
| hyemalis \| aikeni | 7 | 0.843 | 0.500 | 67.2% |
| hyemalis \| caniceps | 138 | 0.687 | 0.745 | 71.6% |
| hyemalis \| dorsalis | 59 | 0.904 | 0.650 | 77.7% |
| dorsalis \| aikeni | 8 | 0.700 | 1.00 | 85.0% |
| dorsalis \| mearnsi | 24 | 0.850 | 0.875 | 86.3% |

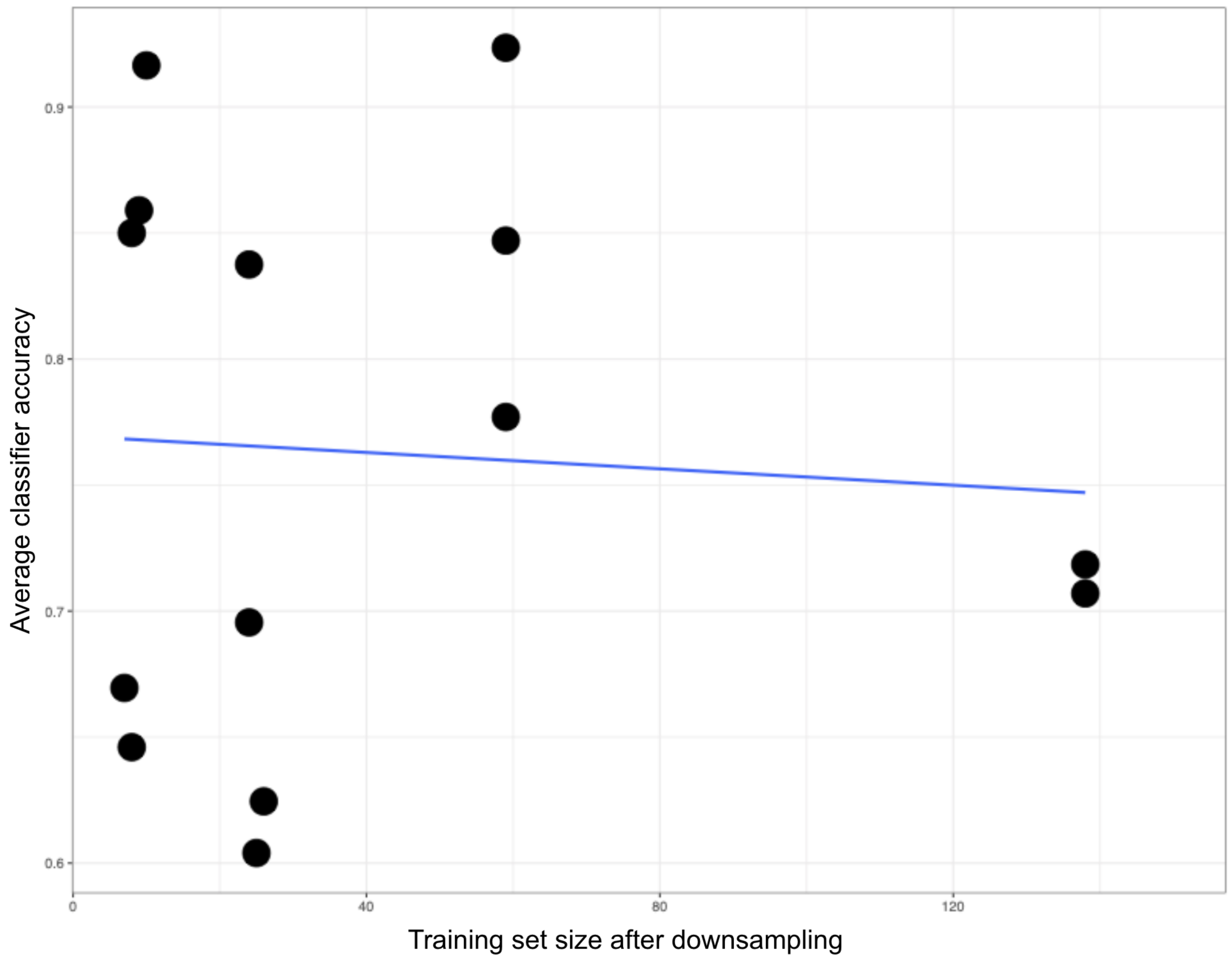


**Figure S12** Random forest classifier's average accuracy for predicting subspecies with different training set sizes. Training set sizes varied because of the amount of songs available for each subspecies. The classifier's accuracy does not appear to be correlated with training set size.

**Table S4** Random forest classifier's variables of importance

| Song feature | Most important | 2nd most important | 3rd most important | Total appearances in top 3 |
|---|---|---|---|---|
| Rate of syllable production | 3 | 3 | 3 | 9 |
| Average syllable lower frequency | 3 | 3 | 1 | 7 |
| Average syllable upper frequency | 3 | 1 | 1 | 5 |
| Average silence duration | 3 | 2 | 0 | 5 |
| Bout duration | 2 | 2 | 1 | 5 |
| Average syllable duration | 1 | 0 | 3 | 4 |
| Number of syllables | 0 | 1 | 3 | 4 |
| Mean syllable stereotypy | 0 | 2 | 2 | 4 |
| Average syllable frequency range | 0 | 1 | 1 | 2 |
| Number of unique syllables | 0 | 0 | 0 | 0 |

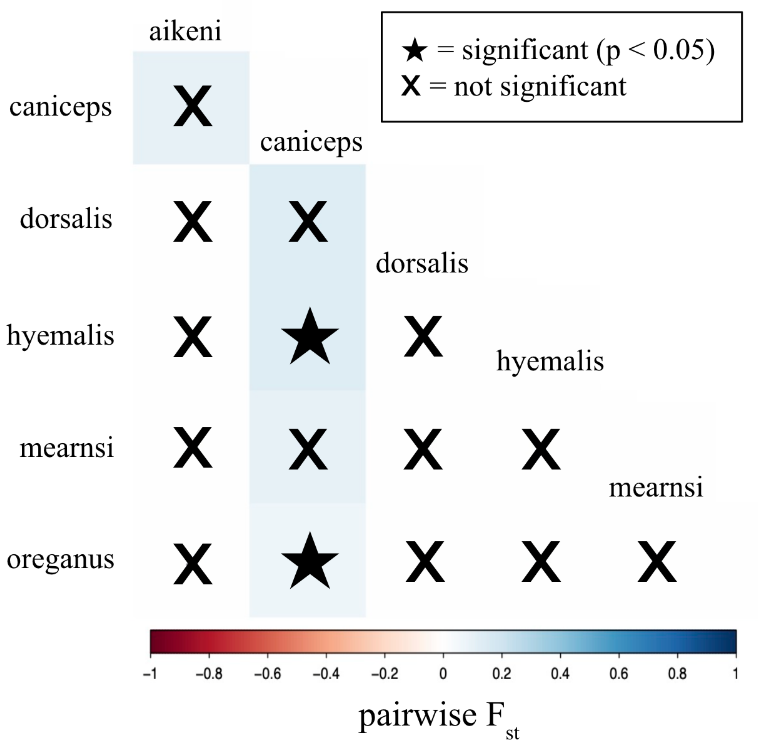


**Figure S13** Pairwise $F_{ST}$ matrix for the COI gene. Subspecies pairs with significant $F_{ST}$ are denoted with stars. The color scale corresponds to $F_{ST}$.

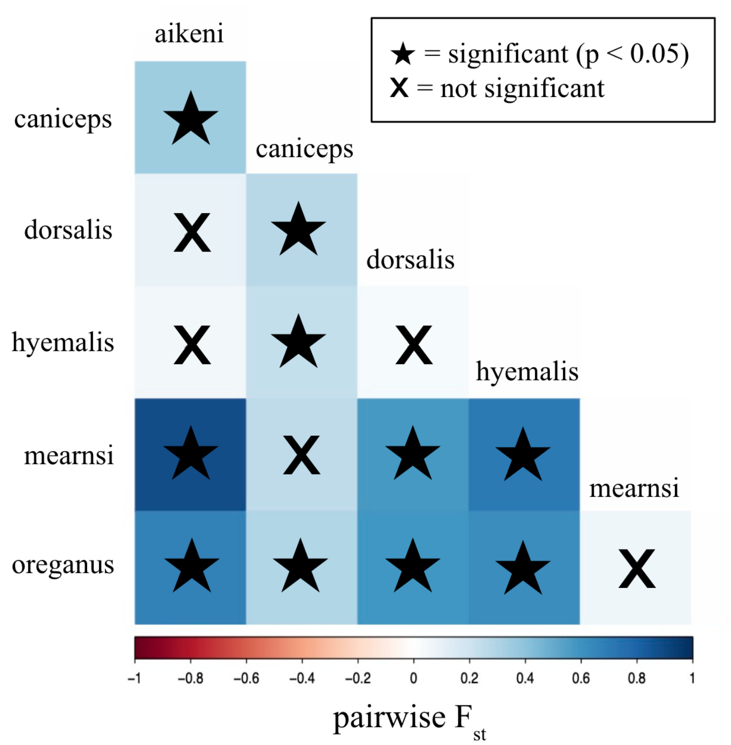


**Figure S14** Pairwise $F_{ST}$ matrix for the ATP gene. Subspecies pairs with significant $F_{ST}$ are denoted with stars. The color scale corresponds to $F_{ST}$.

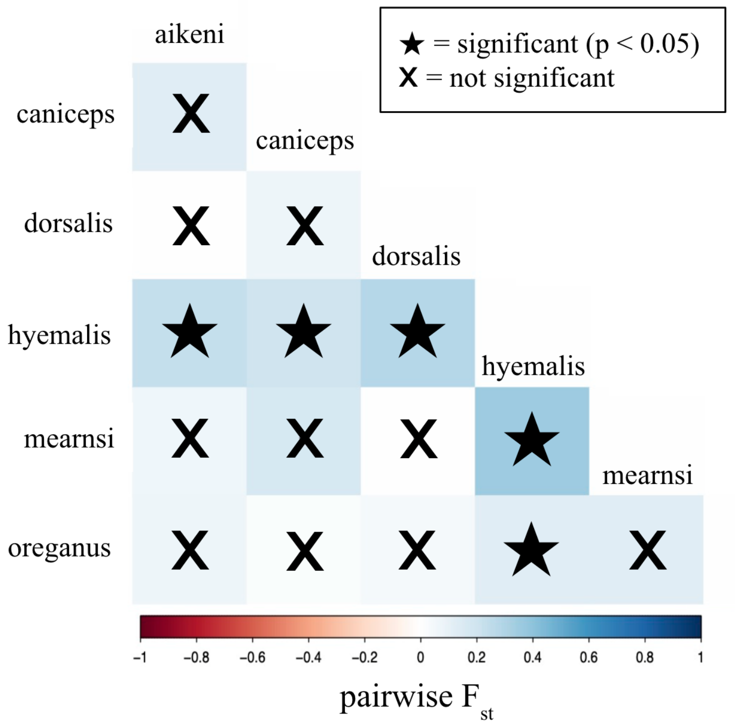


**Figure S15** Pairwise $F_{ST}$ matrix for the FGB gene. Subspecies pairs with significant $F_{ST}$ are denoted with stars. The color scale corresponds to $F_{ST}$.

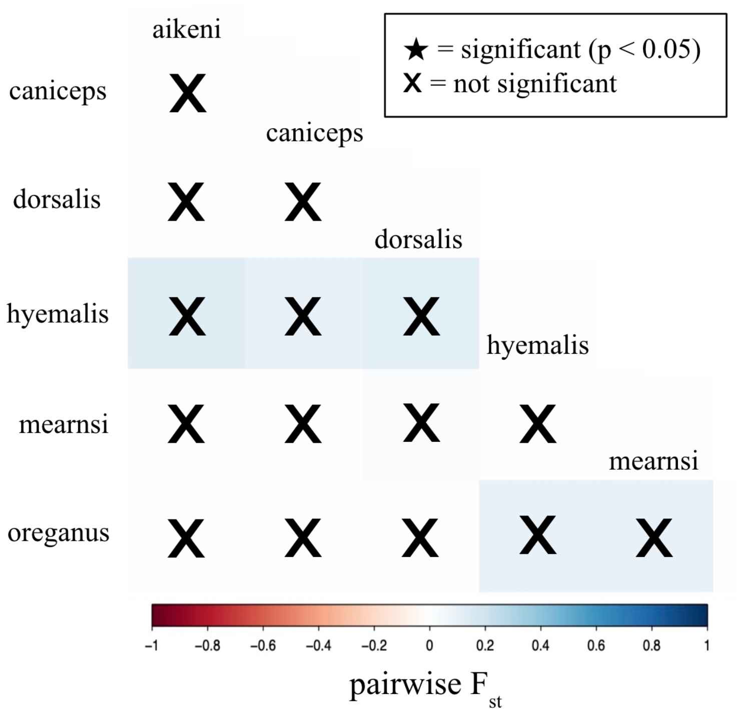


**Figure S16** Pairwise $F_{ST}$ matrix for the ND2 gene. There are no subspecies pairs with significant $F_{ST}$. The color scale corresponds to $F_{ST}$.

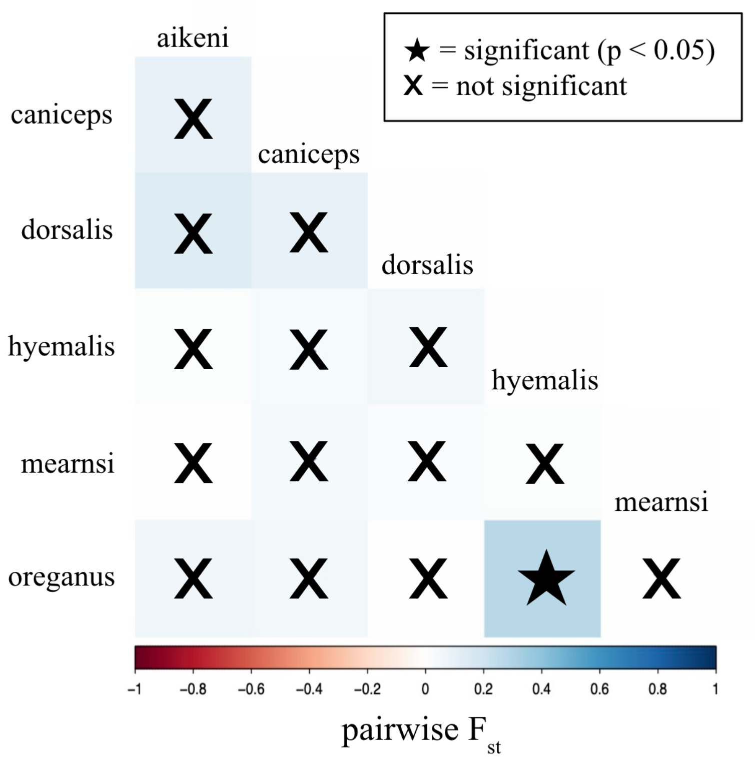


**Figure S17** Pairwise $F_{ST}$ matrix for the CR gene. Subspecies pairs with significant $F_{ST}$ are denoted with stars. The color scale corresponds to $F_{ST}$.

**Table S5** AMOVA for the COI genetic data. 3.2% of the variance was between subspecies and 96.8% of the variance was within subspecies. The genetic data were permuted and an empirical *P*-value of 0.04692 was calculated as the probability that the $F_{ST}$ of the permuted data is greater than the $F_{ST}$ of the observed data.

| **Source of variation** | **Sum of squares** | **Variance components** | **Variation (%)** |
|---|---|---|---|
| Among populations | 1.500 | 0.00635 | 3.229 |
| Within populations | 21.780 | 0.19043 | 96.771 |
| Total | 23.280 | 0.19679 | |

**Table S6** AMOVA for the ATP genetic data. 45.1% of the variance was between subspecies and 54.9% of the variance was within subspecies. The genetic data were permuted and an empirical *P*-value of $P < 1.0 \times 10^{-4}$ was calculated as the probability that the $F_{ST}$ of the permuted data is greater than the $F_{ST}$ of the observed data.

| **Source of variation** | **Sum of squares** | **Variance components** | **Variation (%)** |
|---|---|---|---|
| Among populations | 14.831 | 0.23880 | 45.120 |
| Within populations | 20.913 | 0.29045 | 54.880 |
| Total | 35.744 | 0.52925 | |

**Table S7** AMOVA for the FGB genetic data. 15.2% of the variance was between subspecies and 84.8% of the variance was within subspecies. The genetic data were permuted and an empirical *P*-value of $P < 1.0 \times 10^{-4}$ was calculated as the probability that the $F_{ST}$ of the permuted data is greater than the $F_{ST}$ of the observed data.

| **Source of variation** | **Sum of squares** | **Variance components** | **Variation (%)** |
|---|---|---|---|
| Among populations | 15.458 | 0.14853 | 15.226 |
| Within populations | 79.385 | 0.82693 | 84.774 |
| Total | 94.843 | 0.97546 | |

**Table S8** AMOVA for the ND2 genetic data. 3.0% of the variance was between subspecies and 97.0% of the variance was within subspecies. The genetic data were permuted and an empirical *P*-value of 0.04692 was calculated as the probability that the $F_{ST}$ of the permuted data is greater than the $F_{ST}$ of the observed data.

| Source of variation | Sum of squares | Variance components | Variation (%) |
|---|---|---|---|
| Among populations | 1.866 | 0.00859 | 3.001 |
| Within populations | 19.706 | 0.27754 | 96.999 |
| Total | 21.571 | 0.28613 | |

**Table S9** AMOVA for the CR genetic data. 9.0% of the variance was between subspecies and 91.0% of the variance was within subspecies. The genetic data were permuted and an empirical *P*-value of $P < 1.0 \times 10^{-4}$ was calculated as the probability that the $F_{ST}$ of the permuted data is greater than the $F_{ST}$ of the observed data.

| Source of variation | Sum of squares | Variance components | Variation (%) |
|---|---|---|---|
| Among populations | 27.435 | 0.25556 | 8.998 |
| Within populations | 186.05 | 2.58469 | 91.002 |
| Total | 213.485 | 2.84025 | |

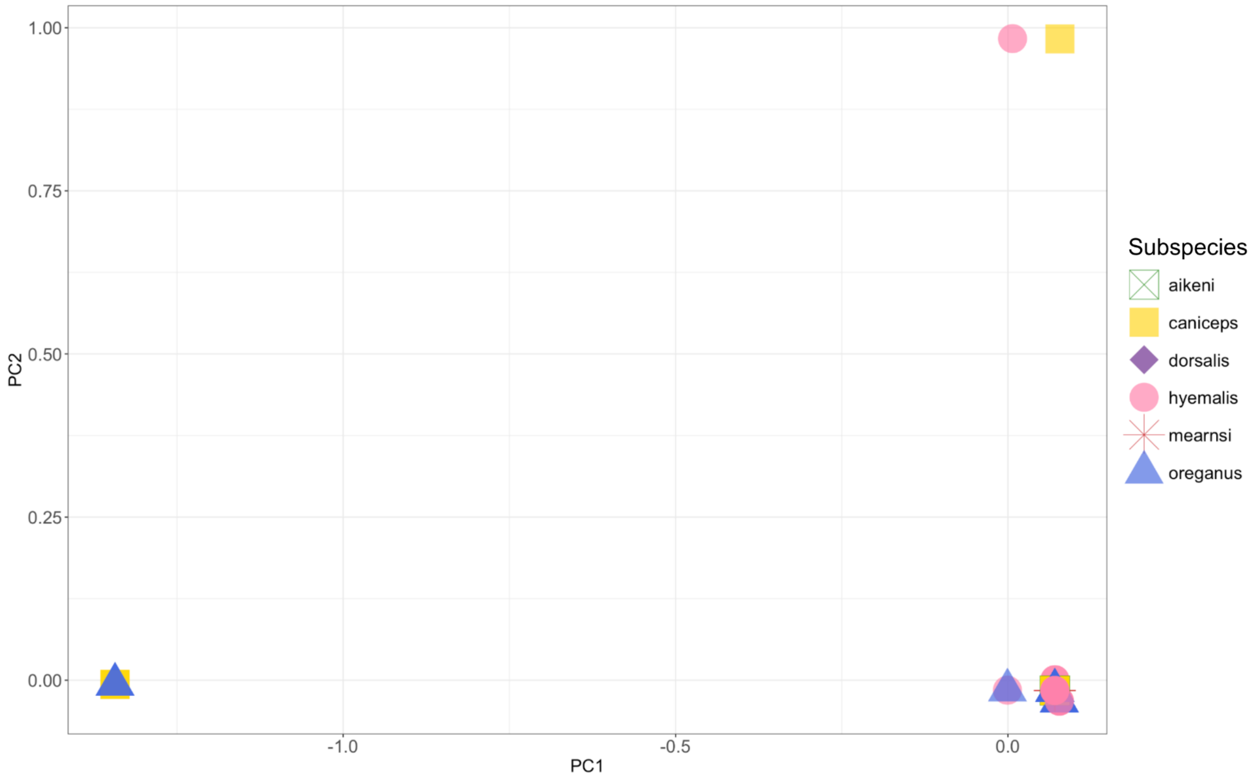


**Figure S18** Principal component analysis results for COI genetic data.

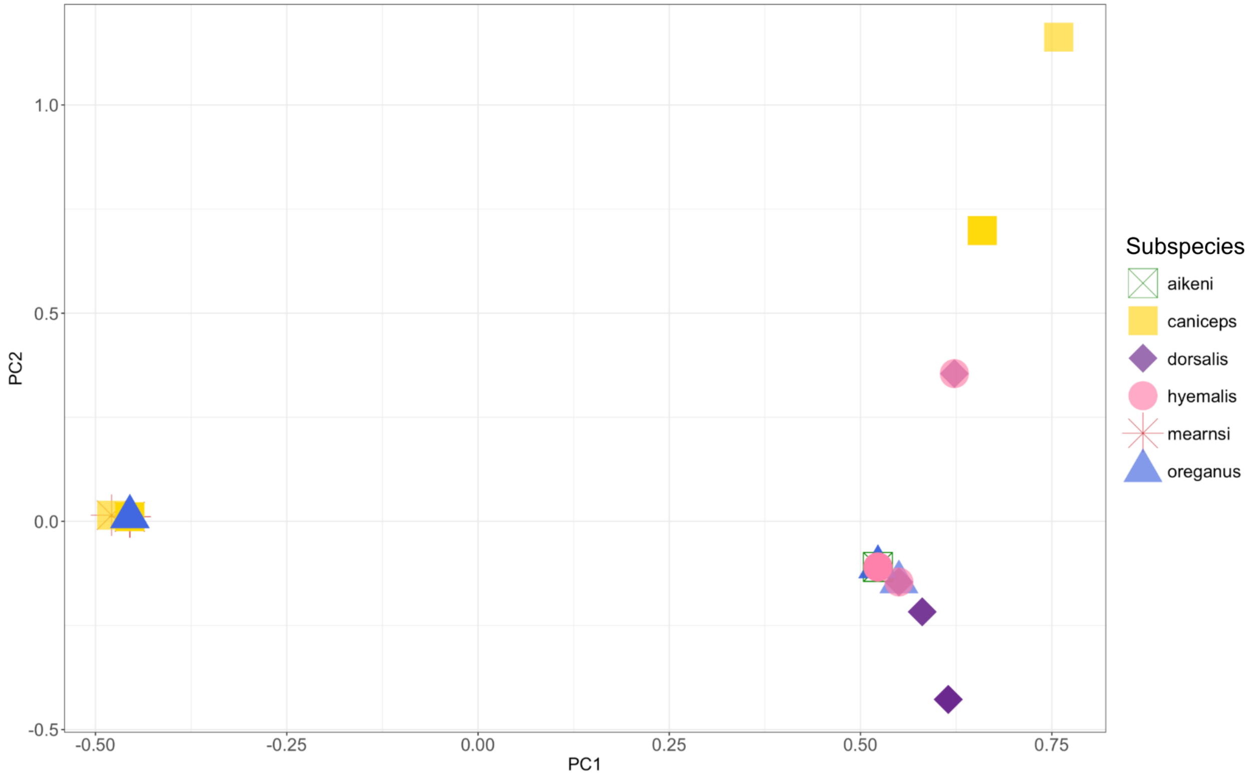


**Figure S19** Principal component analysis results for ATP genetic data.

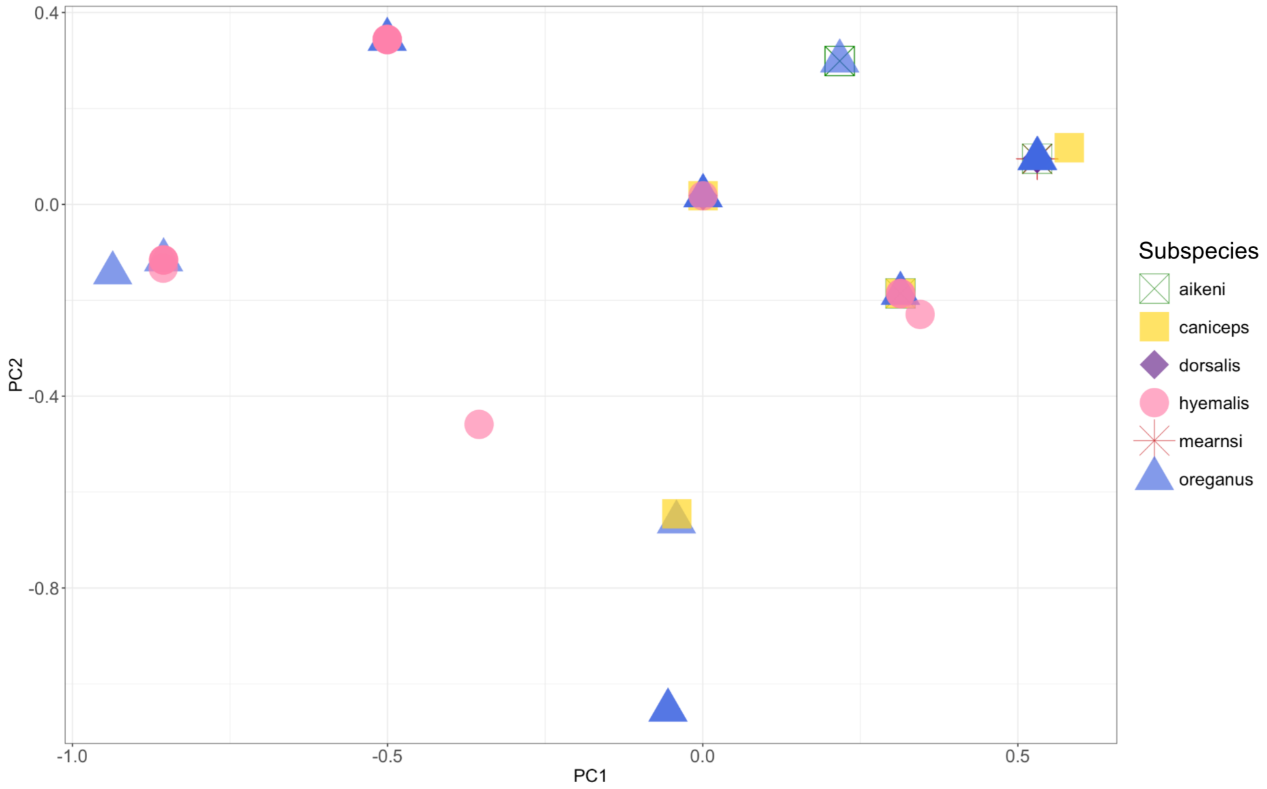


**Figure S20** Principal component analysis results for FGB genetic data.

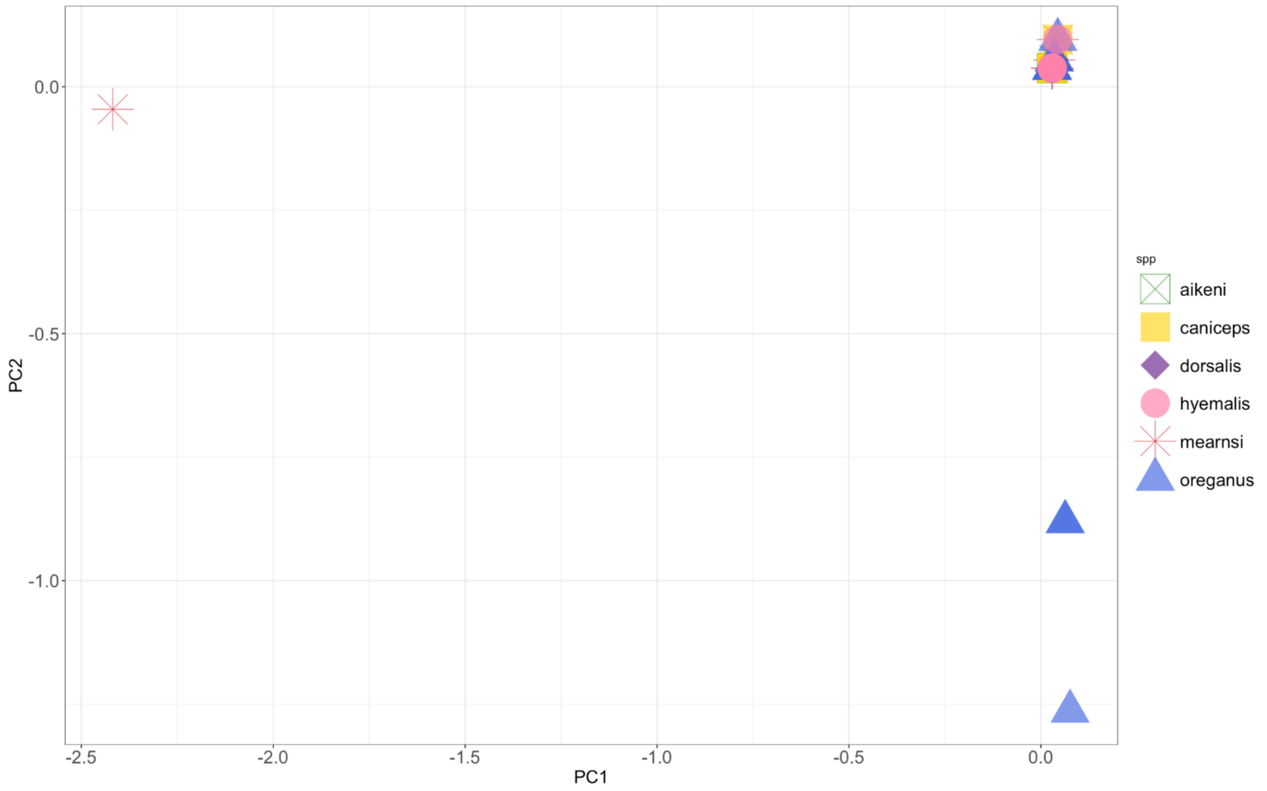


**Figure S21** Principal component analysis results for ND2 genetic data.

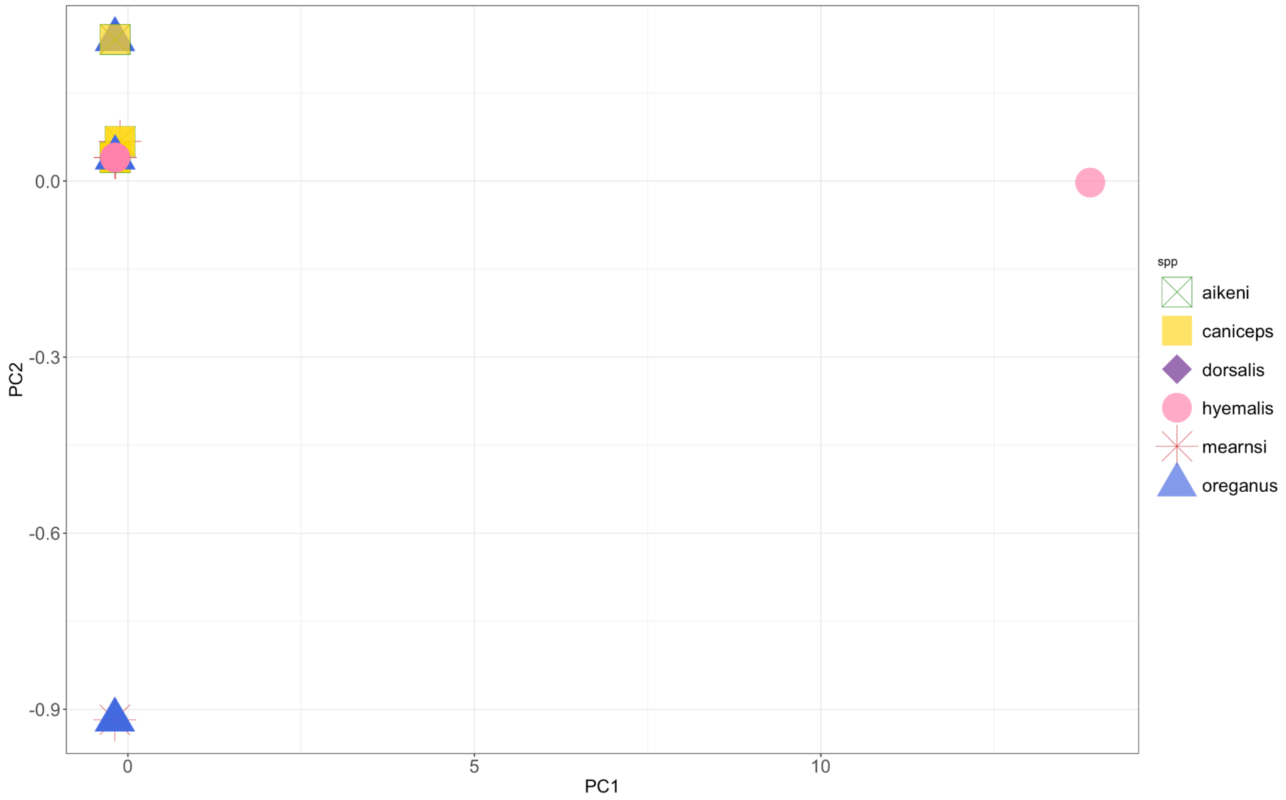


**Figure S22** Principal component analysis results for CR genetic data.

**Table S10** Song PCA areas and proportion of song PCA overlap. The proportion of PCA overlap was calculated by dividing the PCA overlap area by the union of the subspecies' PCA areas (PCA overlap area / (Subspecies 1 PCA area + Subspecies 2 PCA area - PCA overlap area).
Subspecies pair with adjacent ranges that are not known to hybridize
Subspecies pair with hybrid sightings
Subspecies pair without adjacent ranges that are not known to hybridize

| Subspecies Pair | PCA overlap area in PC space | Subspecies 1 PCA area in PC space | Subspecies 2 PCA area in PC space | Percent of PCA overlap |
|---|---|---|---|---|
| dorsalis \| mearnsi | 16.82 | 26.65 | 19.61 | 57.11% |
| dorsalis \| oreganus | 26.62 | 102.6 | 26.65 | 25.94% |
| oreganus \| aikeni | 13.68 | 102.6 | 15.87 | 13.06% |
| hyemalis \| oreganus | 87.79 | 114.5 | 102.6 | 67.88% |
| caniceps \| dorsalis | 24.13 | 36.31 | 26.65 | 62.13% |
| caniceps \| mearnsi | 18.60 | 19.61 | 36.31 | 49.86% |
| caniceps \| oreganus | 34.11 | 102.6 | 36.31 | 32.56% |
| mearnsi \| aikeni | 11.93 | 19.61 | 15.87 | 50.69% |
| mearnsi \| oreganus | 17.89 | 102.6 | 19.61 | 17.15% |
| hyemalis \| aikeni | 14.80 | 114.5 | 15.87 | 12.80% |
| hyemalis \| caniceps | 36.11 | 114.5 | 36.31 | 31.47% |
| hyemalis \| dorsalis | 26.65 | 114.5 | 26.65 | 23.27% |
| hyemalis \| mearnsi | 19.61 | 114.5 | 19.61 | 17.12% |
| dorsalis \| aikeni | 12.13 | 15.87 | 26.65 | 39.90% |
| dorsalis \| mearnsi | 16.82 | 26.65 | 19.61 | 57.11% |
